# EGFR INHIBITION PROMOTES ENTEROENDOCRINE CELL DIFFERENTIATION CONTRIBUTING TO TREATMENT-ASSOCIATED DIARRHEA

**DOI:** 10.64898/2026.06.02.729650

**Authors:** Guilherme Piovezani Ramos, Daniel Zeve, Amy Shepherd, Elisa Saint-Denis, Olivia Alderfer, Bia Frintu, Stanley Dale, Karina Sharma, Emma Mohlmann, Prabhath Mannam, Morgan Byers, Jack Terzian, Claudio Ribeiro, Luiz Fernando Oliveira, Kleiton Silva Borges, Amy E. O’Connell, Diana L. Carlone, Narjust Florez, Meenakshi Rao, David T. Breault

**Author notes:** Corresponding Author: David Breault, MD, PhD Division of Endocrinology, Department of Pediatrics, Boston Children’s Hospital, Harvard Medical School 300 Longwood Avenue, Boston, MA 02115, USA. Equal contribution (Guilherme Piovezani Ramos, Daniel Zeve).

## Abstract

Enteroendocrine cells (EECs) are specialized sensors of the gastrointestinal (GI) epithelium that regulate gut function and systemic metabolism through hormone secretion. The molecular pathways directing intestinal stem cell (ISC) differentiation into EECs are incompletely understood due, in part, to their rarity. We sought to identify novel regulators of human EEC differentiation using a high-throughput screen of FDA-approved drugs and human duodenal organoids. Two epidermal growth factor receptor inhibitors (EGFRi) commonly used in cancer therapy and known to cause GI side effects, erlotinib and lapatinib, emerged as strong inducers of EEC differentiation, dramatically increasing chromogranin A (CHGA) expression compared to controls, while maintaining ISC function and organoid growth. EGFRi-treated organoids revealed robust and broad upregulation of EEC hormones, including serotonin (5HT), motilin (MLN), and somatostatin (SST), among others. In agreement with these findings, analysis of a patient cohort with lung cancer revealed an association with erlotinib use and increased circulating levels of the above EEC hormones compared to matched controls. Supporting a direct effect of EGFRi on EEC differentiation, mice treated with erlotinib demonstrated increased EEC numbers and hormones and showed EGFRi-associated diarrhea (EAD), a limiting side effect of these medications. Mechanistically, EGFRi induced upregulation of interferon (IFN) signaling targets during early ISC-to-EEC differentiation. Consistent with this, inhibition of Signal Transducer and Activator of Transcription 1 (STAT1) attenuated EGFRi-induced EEC differentiation. These findings provide important insight into EEC differentiation that could inform treatment strategies for EAD, metabolic diseases, and GI diseases.

**Brief Summary:** Inhibition of EGFR signaling promotes human ISC-to-EEC differentiation through activation of STAT1 signaling.

## INTRODUCTION

The intestinal epithelium is maintained through the continual activity of intestinal stem cells (ISCs) and absorptive and secretory progenitor cells, which give rise to differentiated cell types, including enterocytes, goblet cells, Paneth cells, tuft cells, and enteroendocrine cells (EECs)(*1*). Ongoing efforts focus on defining the signaling pathways underlying differentiation of these cell types, which relies on the interplay of intrinsic transcriptional programs and extrinsic signaling gradients along the crypt-villus axis(*2–4*). These gradients and programs are highly context-dependent and can differ between species, highlighting the importance of robust human-centered models such as ISC-derived intestinal organoids(*5, 6*).

EECs comprise approximately 1% of the gastrointestinal (GI) epithelium, making them one of the rarest cell populations within the intestine(*7*). Despite their rarity, EECs represent the largest endocrine system in the human body, both by cell number and hormonal diversity(*8*). In health, EECs respond to intestinal luminal contents by releasing hormones important for regulation of a variety of key functions, including GI motility, pancreatic function and appetite control(*8*). Individual EECs secrete a range of hormones, including serotonin (5HT), motilin (MLN), somatostatin (SST), glucose-dependent insulinotropic polypeptide (GIP), ghrelin (GHRL), glucagon-like peptide 1 (GLP-1), peptide YY (PYY), and cholecystokinin (CCK), depending on their location in the GI tract. In disease, changes in EEC number or activity have been associated with GI disorders (irritable bowel syndrome, inflammatory bowel disease), and metabolic disorders (obesity, type II diabetes mellitus), as illustrated by the groundbreaking impact of GIP/GLP-1 receptor agonists(*9*). Further understanding of the factors governing EEC differentiation may lead to the development of novel therapeutics for GI and metabolic diseases.

Compared to other mature epithelial cell types, progress in understanding the factors that drive EEC differentiation has been technically challenging, in part hampered by their rarity and diversity(*10–12*). Efforts to understand the mechanisms controlling EEC differentiation from ISCs have focused on the role of specific signaling pathways and key transcription factors. For example, previous studies have shown that withdrawal of wingless-related integration site (WNT) and epidermal growth factor (EGF) favors secretory lineage specification at the cost of reduced organoid growth and forced quiescence of ISCs(*10, 13*). In addition, studies in murine intestinal organoids showed that inhibiting the EGF receptor (EGFR), either in the presence or absence of EGF, is not sufficient to promote EEC differentiation(*10*). In contrast, overexpression of NEUROG3 exclusively drives EEC differentiation at the expense of the other intestinal cell lineages(*11, 12*). Finally, the addition of small molecule inhibitors to a base differentiation media, targeting a combination of endocannabinoid, JNK, and/or FOXO1 signaling, promotes robust EEC differentiation while maintaining long-lived, multipotent ISCs(*14*).

Here, we performed a small molecule screen with human duodenal organoids, which identified two EGF receptor inhibitors (EGFRi)—erlotinib and lapatinib, commonly used in cancer therapy and known to cause gastrointestinal (GI) toxicity including diarrhea—as potent inducers of EEC differentiation while maintaining self-renewing ISCs and multilineage differentiation. EGFRi-treated human organoids demonstrated increased expression of EEC-specific markers, including *TPH1* (encoding tryptophane hydroxylase required for generating 5HT), *MLN,* and *SST*, among others. Analysis of a patient cohort with lung cancer revealed an association between erlotinib use and increased circulating levels of the above EEC hormones compared to age, sex, and disease-matched controls. Confirming a direct effect of EGFRi on EEC differentiation, mice treated with erlotinib demonstrated increased EEC numbers and hormones and showed EGFRi-associated diarrhea (EAD). Mechanistic studies revealed EGFRi promotes EEC differentiation through upregulation of Signal Transducer and Activator of Transcription 1 (STAT1) and Interferon (IFN) target genes. Collectively, these data establish a link between EGFR-STAT1 signaling and EEC differentiation, with direct translational implications for understanding EAD, a therapy-limiting side effect of EGFR inhibitors.

## RESULTS

### EGFR inhibitors promote ISC-to-EEC differentiation in human duodenal organoids

To elucidate factors that promote EEC differentiation, we conducted a high-throughput screen of over 600 FDA-approved drugs with human duodenal organoids (**Supplementary Table 1**). Organoids were cultured in base differentiation media(*14*) for nine days (which served as a negative control) or with the addition of each compound, in duplicate, and assessed for CHGA expression using high-speed confocal immunofluorescence (IF) imaging (**Fig 1A**). As a positive control, organoids were treated with the FOXO1 inhibitor AS1842856, a known inducer of EEC differentiation(*14*). At the end of the differentiation period, CHGA staining identified erlotinib, a first-generation EGFR inhibitor (inhibits EGFR/ErbB1), and lapatinib, a second-generation EGFR inhibitor (inhibits EGFR/ErbB1, ErbB2, ErbB4), as potent inducers of EEC differentiation, having comparable levels of staining to the positive control (**Fig 1B, Supplementary Fig 1**), highlighting a potential role for EGFR family members in regulating EEC differentiation.

**Figure 1.**
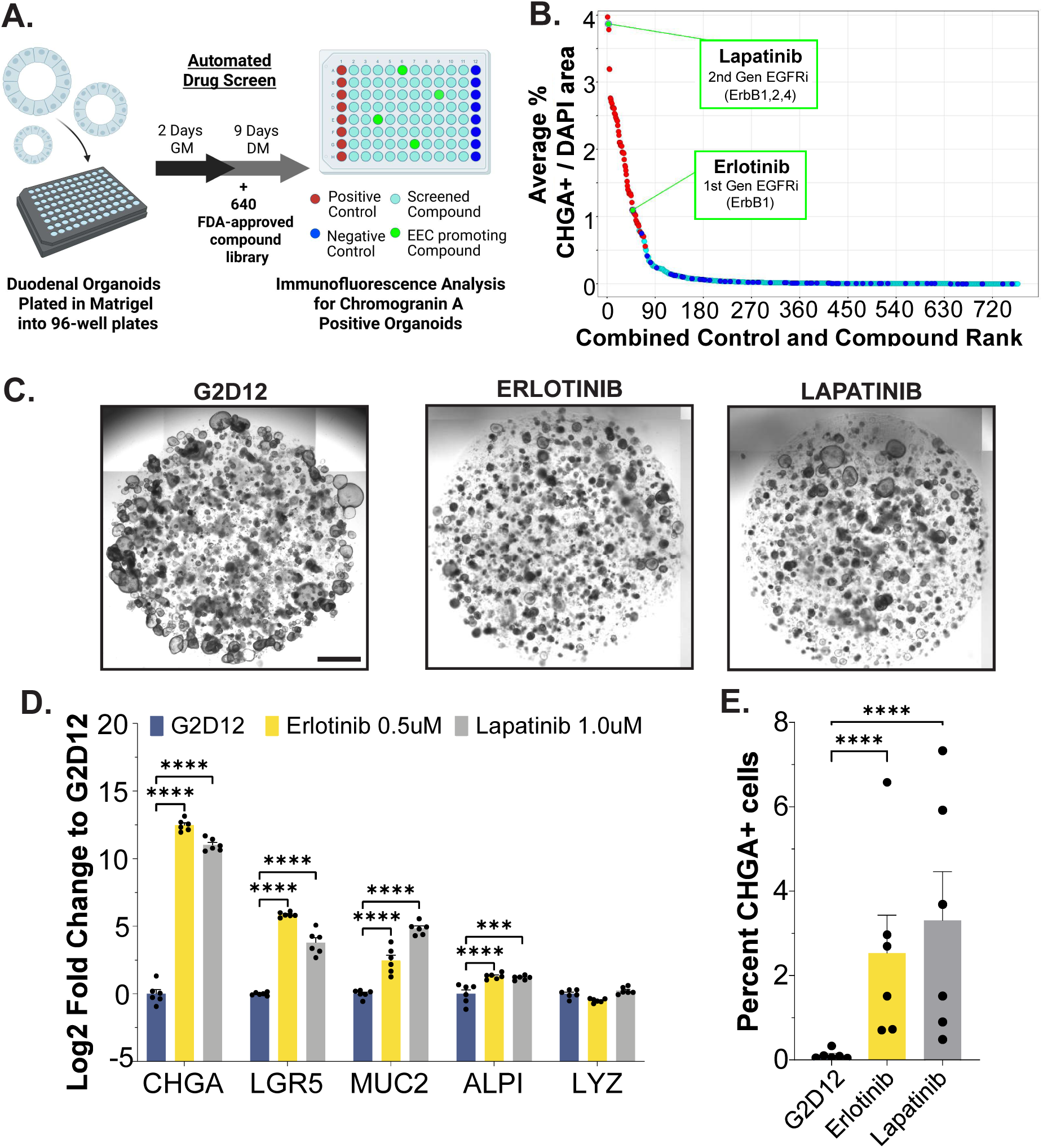
High-throughput Screening of FDA-approved Small Molecules Identifies EGFR Inhibitors as Inducers of Human Enteroendocrine Cell Differentiation. (A) Schematic representation of the high-throughput screen in a 96-well plate format, highlighting the use of human duodenal organoids and our differentiation protocol involving two days of growth media (GM) followed by nine days of differentiation media (DM, with and without small molecules). (B) Ranking of compounds based on average percentage of CHGA⁺ fluorescence signal relative to total DAPI area. Red denotes positive control (AS-treated organoids), dark blue represents negative control (DM alone), light blue represents screened compounds, and green represents compounds that induce EEC differentiation, including erlotinib and lapatinib. (C) Representative brightfield microscopy of human duodenal organoids (whole well) grown for two days in GM followed by 12 days in DM (G2D12), G2D12 with erlotinib, and G2D12 with lapatinib. Scale bar = 1 mm. (D) qPCR analysis of intestinal lineage markers in organoids grown in G2D12, G2D12 with erlotinib, and G2D12 with lapatinib, normalized to ACTB and plotted as log₂ fold change relative to G2D12. Representative experiment showing n = 6 wells per condition from a single organoid line. CHGA = chromogranin A; LGR5 = leucine-rich repeat-containing G-protein coupled receptor 5; MUC2 = mucin 2; ALPI = intestinal alkaline phosphatase; LYZ = lysozyme. (E) Flow cytometry analysis of CHGA⁺ cells from organoids grown in G2D12, G2D12 with erlotinib, and G2D12 with lapatinib. Data represent averages from six independent experiments using six different organoid lines. Bars show mean ± SEM; two-way ANOVA with Tukey correction for multiple comparisons (D); one-way ANOVA with Tukey correction for multiple comparisons (E). ***p < 0.001; ****p < 0.0001. Each experiment was repeated with at least three independent organoid lines.

To validate the screening results, we performed a dose-response analysis to establish the optimal concentrations of erlotinib and lapatinib required to induce EEC differentiation (**Supplementary Fig 2A-B**). For these studies, human organoids were first cultured for two days in growth media to expand the ISC population and then cultured in base differentiation media for 12 days (G2D12) (**Fig 1C**). Addition of erlotinib (0.5μM) or lapatinib (1μM) led to markedly increased induction in *CHGA* expression, compared to untreated G2D12 controls, measured by real time quantitative PCR (qPCR) (**Fig 1D**). EGFRi treatment also induced a significant, but less robust, increase in expression of other intestinal lineage markers, including *LGR5* (stem cells), *MUC2* (goblet cells), and *ALPI* (enterocytes), but not *LYZ* (Paneth cells). Similar to our previous study demonstrating that small molecule-induced EEC differentiation is accompanied by broader changes in intestinal epithelial lineage composition, these findings suggest that EGFRi promotes a general shift toward a more differentiated epithelial state while preferentially expanding the secretory lineage(*14*). Consistent with this, flow cytometric analysis confirmed significantly higher numbers of CHGA+ cells following treatment with erlotinib (2.53 ± 0.89%) or lapatinib (3.30 ± 1.15%), greater than 20 times higher than controls (0.11 ± 0.04%) (**Fig 1E, Supplementary Fig 2C**). Given our primary interest in the factors governing EEC differentiation and the marked induction of EEC-specific markers and hormone production observed, subsequent analyses focused on EEC biology. Together, these data establish that inhibition of EGFRs promotes human EEC differentiation in organoids.

### EGFRi leads to increased production of EEC hormones in human organoids

Hormonal expression within EECs varies depending on location within the GI tract, with individual EECs typically producing one to two specific hormones. Spatial and cellular specialization is further reflected at the transcriptional level by distinct lineage branch points that bias differentiation toward 5HT-producing or peptide hormone-producing cells(*9*). To characterize the hormones induced by erlotinib or lapatinib, we first assessed gene expression in duodenal organoids differentiated for 12 days (**Fig 2A**). qPCR demonstrated significant induction of *TPH1*, *MLN, SST, GIP,* and *GHRL* compared to controls (**Fig 2B**). To verify these findings, we performed IF, which revealed CHGA, 5HT, and GIP expression in >60% of treated organoids compared to <5% in controls (**Fig 2C-G, Supplementary Fig 3A-B**). Finally, we performed ELISA to detect secreted hormones following forskolin stimulation, which revealed marked induction of both 5HT and GIP following treatment with EGFRi (**Fig 2H-I**). Together, these results show that EGFRi promotes EEC differentiation into both 5HT-producing and peptide hormone-producing EECs commonly located in the human duodenum.

**Figure 2.**
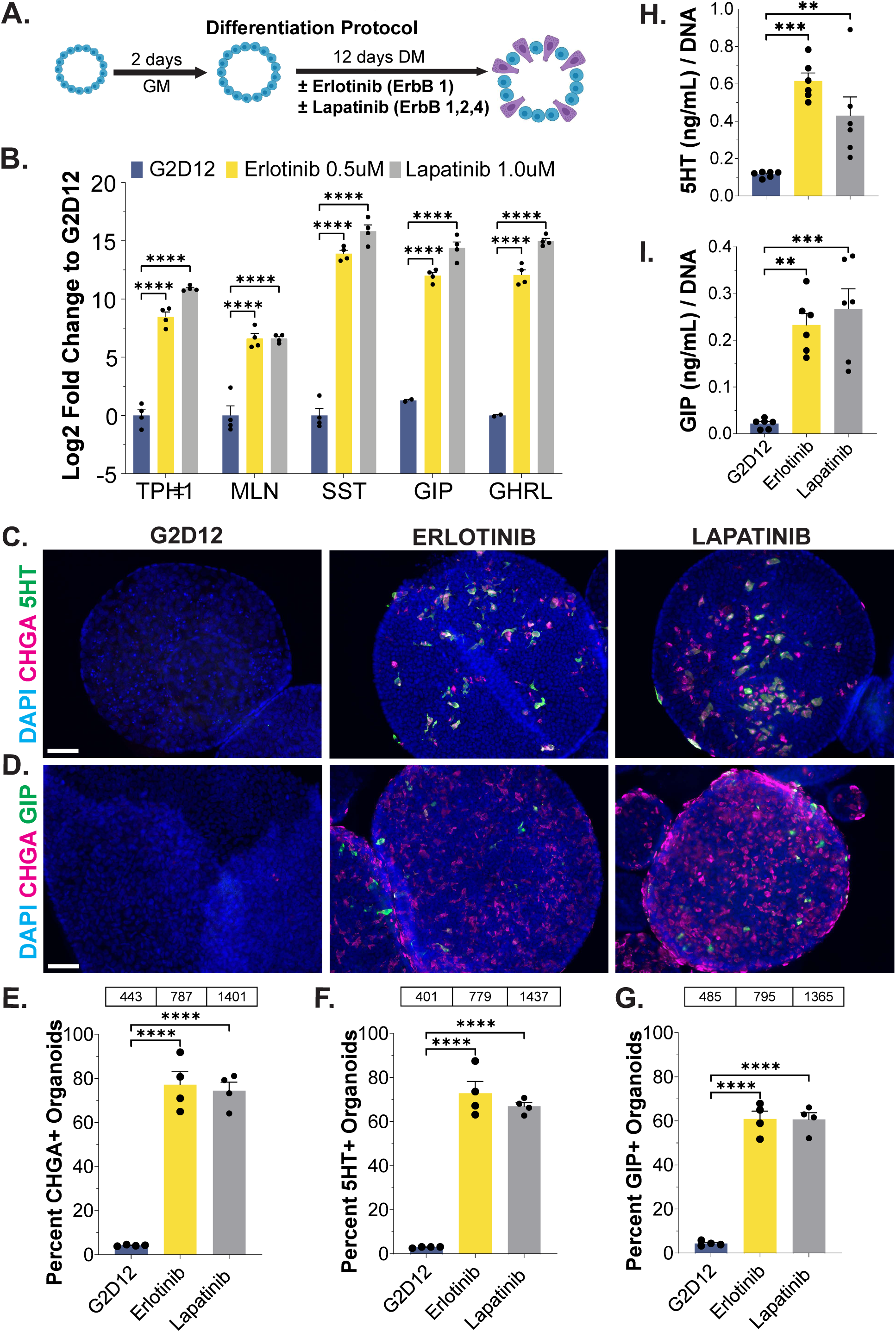
Differentiation with Erlotinib and Lapatinib Induces EEC Hormone Production and Secretion. (A) Schematic of our differentiation protocol, in which human duodenal organoids are grown for two days in growth media (GM) followed by 12 days in differentiation media (DM) with and without erlotinib or lapatinib. (B) qPCR analysis of EEC markers in organoids grown in G2D12, G2D12 with erlotinib, and G2D12 with lapatinib, normalized to ACTB and plotted as log₂ fold change relative to G2D12. Representative experiment showing n = 2–4 wells per condition from a single organoid line. TPH1 = tryptophan hydroxylase 1; MLN = motilin; SST = somatostatin; GIP = glucose-dependent insulinotropic peptide; GHRL = ghrelin. (C,D) Representative immunofluorescence staining of CHGA (magenta) with 5HT (green, (C)) or GIP (green, (D)) in organoids grown in G2D12, G2D12 with erlotinib, and G2D12 with lapatinib. Nuclei (DAPI, blue). Scale bars = 50 µm. (E–G) Percentage of organoids positive for CHGA (E), 5HT (F), and GIP (G) in G2D12, G2D12 with erlotinib, and G2D12 with lapatinib. Tables above graphs indicate the total number of organoids analyzed per condition. Data represent averages from four independent experiments using four different organoid lines. (H) 5HT ELISA of conditioned media from organoids grown in G2D12, G2D12 with erlotinib, and G2D12 with lapatinib, collected after 24 hours of forskolin stimulation on day 14. Data are normalized to total DNA per sample. Representative experiment showing n = 6 wells per condition from a single organoid line. (I) GIP ELISA of conditioned media from organoids grown in G2D12, G2D12 with erlotinib, and G2D12 with lapatinib, collected after 24 hours of forskolin stimulation on day 14. Data are normalized to total DNA per sample. Representative experiment showing n = 6 wells per condition from a single organoid line. Bars show mean ± SEM; two-way ANOVA with Tukey correction for multiple comparisons (B); one-way ANOVA with Tukey correction for multiple comparisons (E–I). **p < 0.01; ***p < 0.001; ****p < 0.0001. Each experiment was repeated with at least three independent organoid lines.

### Erlotinib treatment is associated with increased serum levels of EEC hormones in humans

While EGFRi therapy is central to the treatment of a variety of malignancies, including lung cancer, their use is frequently limited by GI toxicity(*15*). EGFRi-associated diarrhea (EAD) is a significant side effect impacting 30–90% of patients treated with this class of drugs(*16*). Although several mechanisms have been proposed, the precise pathophysiology underlying EAD remains enigmatic^10-12^. Given the above results, we hypothesized that changes in EEC hormones may account, at least in part, for EAD, as EEC hormones play roles in both gut motility and fluid absorption and secretion. To assess this, we sought to measure serum levels of EEC hormones in patients with lung cancer treated with EGFRi. We identified a cohort of 94 patients with lung adenocarcinoma, 47 treated with erlotinib and 47 gender- and age-matched controls treated with non-EGFRi based chemotherapy (**Fig 3A**, **Supplementary Table 2**). Analysis of EEC hormone levels in serum obtained as part of routine clinical care revealed significantly higher levels of 5HT, MLN, and SST — hormones selected for their relative stability in serum compared to other EEC hormones subject to rapid enzymatic degradation — in erlotinib-treated patients compared to their matched controls (**Fig 3B-C**). This association raises the intriguing possibility that changes in EECs numbers and/or their hormones in response to EGFRi treatment may contribute to EAD, motivating direct causal investigation in a controlled model system.

**Figure 3.**
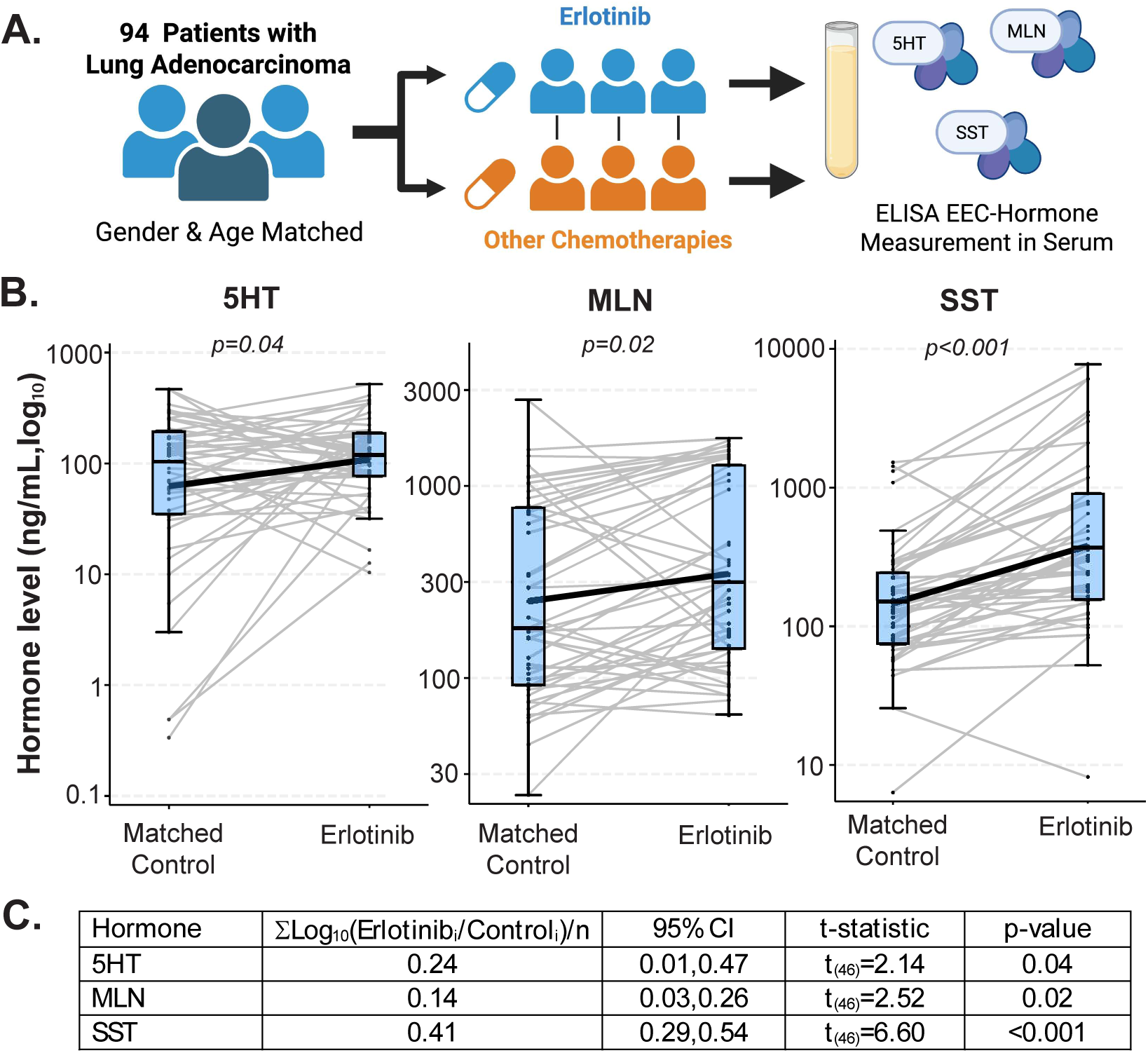
Erlotinib Treatment Is Associated with Increased Circulating EEC Hormones in Patients with Lung Adenocarcinoma. (A) Schematic of study design. Serum samples were obtained from patients with lung adenocarcinoma (n = 94 total; 47 erlotinib-treated and 47 age- and gender-matched controls treated with non–EGFR inhibitor-based chemotherapy). Circulating EEC hormones were measured by ELISA. (B) Serum levels of 5HT, MLN, and SST in matched control and erlotinib-treated patients. Each line represents a matched pair. Hormone concentrations in ng/mL are plotted on a log₁₀ scale to account for variability in circulating hormone levels and differences in baseline concentration ranges between hormones. (C) Summary of log₁₀(erlotinib/control, n=47) fold change for each matched pair, with 95% confidence intervals, t-statistics, and corresponding p-values. Fold-change analysis was performed to directly compare hormone levels between matched control and erlotinib-treated patients while accounting for inter-patient variability. Statistical significance was determined using paired, two-tailed Student’s t-test. p-values are indicated in (B) and (C).

### Erlotinib promotes EEC differentiation, hormone production, and diarrhea in mice

Given the associations between erlotinib use and elevated serum hormone levels, we next sought to determine whether a causal link exists between erlotinib use and changes in EECs, hormone levels and/or GI function, *in vivo*. Treatment of C57BL/6J mice with erlotinib (120 mg/kg/d for 7 days(*17*)) (**Fig 4A**) resulted in significant weight loss beginning at 3 days compared to vehicle-treated controls (**Fig 4B**). In addition, nearly 80% of erlotinib-treated mice developed diarrhea, as confirmed by increased percentage of stool water content (**Fig 4C-D**). This occurred despite the mice eating and drinking less than the vehicle-treated controls, indicating a dramatic dysregulation of water absorption (**Fig 4E-F**). To assess gross changes in the intestine that could account for these findings, we performed histological analysis, which revealed no significant changes in crypt-villus architecture or inflammatory infiltrates (**Fig 4G-H, Supplementary Fig 4A-B**). In contrast, and consistent with the effect of EGFRi on EEC differentiation, the duodenum of erlotinib-treated animals demonstrated a significant increase in CHGA+ cells compared to controls (**Fig 4I**). Further analysis of the duodenum from erlotinib-treated mice confirmed increased expression of the EEC markers *Chga*, *Tph1*, *Sst, and Gip*, as well as increased expression of the other intestinal epithelial markers, including *Lgr5* and *Muc2* (**Fig 4J, Supplementary Fig 4C**), which closely mirrored our human organoid findings (**Fig 1D and 2B**). Next, we quantified circulating levels of EEC hormones in the serum, which revealed significant elevations in 5HT, GLP-1, and PYY in erlotinib-treated mice, without changes in SST, GHRL, or GIP (**Fig 4K, Supplementary Fig 4D**). Together, these data confirm the findings from the human organoid and serum studies and indicate that EGFRi treatment of mice increased EEC numbers and hormone production, in association with watery diarrhea.

**Figure 4.**
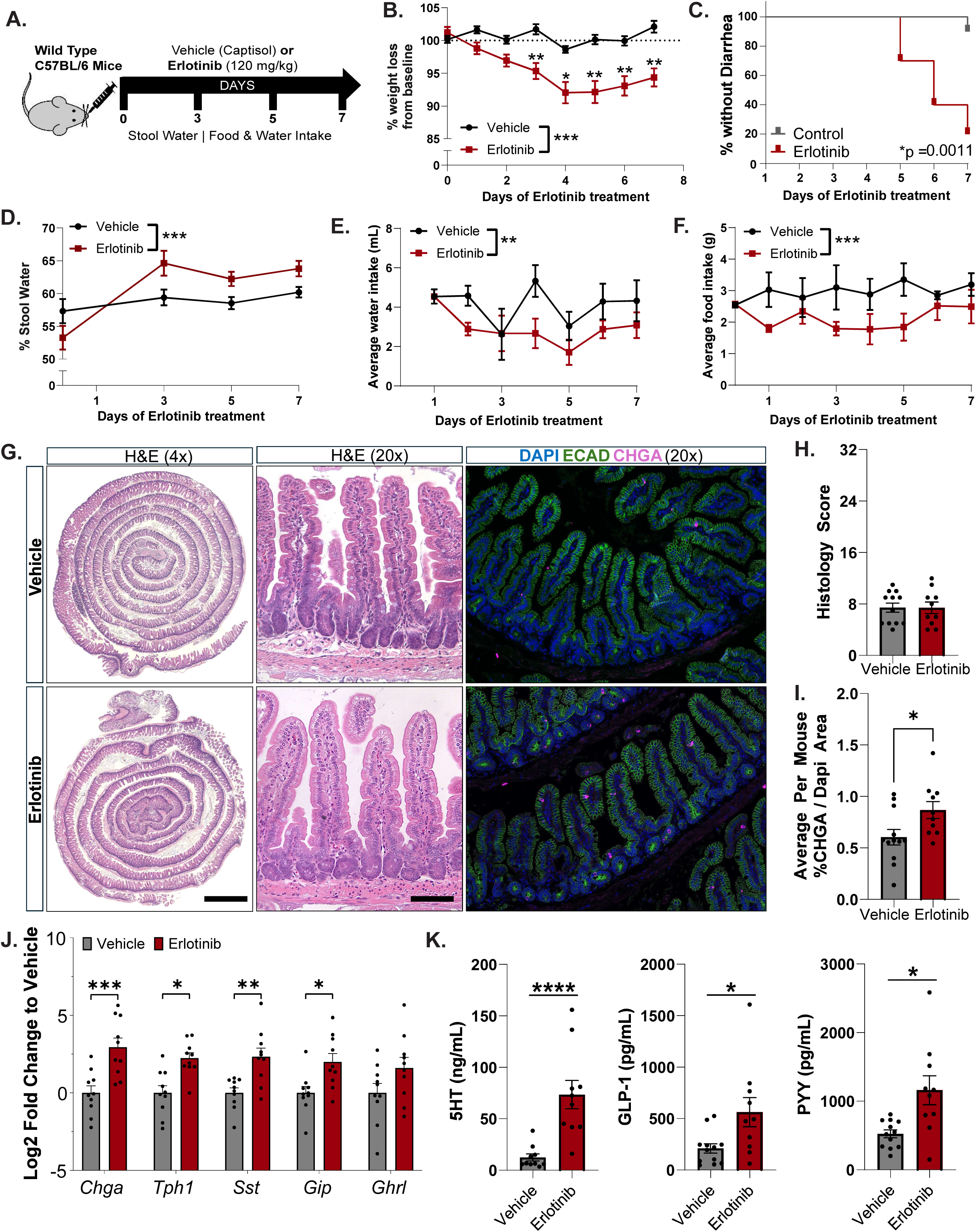
Erlotinib Treatment Promotes EEC Differentiation, Hormone Production, and Diarrhea in Mice. (A) Schematic of our *in vivo* treatment protocol. Wild-type C57BL/6 mice (n = 10 per group) were treated daily with vehicle (Captisol) or erlotinib (120 mg/kg) for 7 days, with serial assessment of body weight, stool output, food intake, and water intake. (B) Percent weight change from baseline over time in vehicle- and erlotinib-treated mice. (C) Kaplan–Meier analysis of time to development of diarrhea, defined by stool scoring criteria, during treatment. (D) Stool water percentage, calculated from wet and dry stool weights following pellet collection and drying. (E,F) Average water intake (E) and food intake (F) per mouse, measured daily and normalized by animals per cage. (G) Representative duodenal histology from vehicle- and erlotinib-treated mice, including H&E staining (4x and 20x) and immunofluorescence for CHGA (magenta), E-cadherin (green), and nuclei (DAPI, blue). Scale bars = 1 mm (4×) and 100 µm (20×). (H) Histological scoring of epithelial damage and inflammatory infiltrate. (I) Quantification of CHGA+ cells in duodenal epithelium, measured as CHGA+ pixel area within the E-cadherin–defined epithelial compartment and normalized to DAPI area. (J) qPCR analysis of intestinal epithelial and EEC markers in duodenal tissue, normalized to Actb and plotted as log₂ fold change relative to vehicle-treated mice. Representative experiment from one cohort of mice. Chga = chromogranin A; Tph1 = tryptophan hydroxylase 1; Sst = somatostatin; Gip = glucose-dependent insulinotropic peptide; Ghrl = ghrelin. (K) Circulating levels of EEC hormones (5HT, GLP-1, and PYY) measured by ELISA from plasma or serum collected on day 7. Bars show mean ± SEM; mixed-effects analysis with Tukey correction for multiple comparisons (B, D–F); two-way ANOVA with Tukey correction for multiple comparisons (J); Welch’s t-test (H, I, K); log-rank (Mantel–Cox) test for Kaplan–Meier analysis (C). *p < 0.05; **p < 0.01; ***p < 0.001; ****p < 0.0001. Experiments were repeated in three independent cohorts of mice.

### EGFRi-induced STAT1 signaling drive ISC-to-EEC differentiation

To define the molecular mechanisms underlying EGFRi–mediated differentiation, we examined early signaling events during ISC-to-EEC differentiation in human duodenal organoids. Time-course analysis revealed upregulation of *CHGA* as early as day 4 following lapatinib treatment, which preceded the induction of *TPH1* on day 6 (**Supplementary Fig 5A-B**). To identify changes prior to *CHGA* expression, we performed bulk RNA-sequencing 48 hours after treatment with lapatinib (G2D2+lapa). Organoids cultured in standard differentiation media (G2D2) served as the negative control, while AS1842856 (AS) was employed as a positive control for EEC induction (G2D2+AS) (**Fig. 5A**). Analysis of differentially expressed genes (DEGs) identified 5,175 transcripts between lapatinib and G2D2 and 8,761 between AS and G2D2 (**Supplementary Fig 5C**). When comparing the two treatments, normalized to the negative control, overlapping genes segregated into five clusters, with two showing coordinated upregulation (Cluster 4) or downregulation (Cluster 2) in response to both lapatinib and AS (**Fig. 5B**). Further analysis of Cluster 4 revealed DEGs enriched for inflammatory and antiviral pathways, including those involved in IFN signaling (**Fig 5C, Supplementary Fig 5D**). Gene set enrichment analysis (GSEA) of lapatinib-induced DEGs further revealed significant enrichment of genes involved in IFN response (**Fig. 5D, Supplementary Fig 5E**). Consistent with a response to lapatinib treatment, analysis of downregulated DEG in Cluster 2 revealed suppression of canonical EGFR-associated pathways, including mTORC signaling (**Fig 5E**, **Supplementary Fig 5F-G**). We next used ingenuity pathway analysis (IPA) to identify putative mediators of the EGFRi-induced response, which highlighted IRF1 and STAT1 as potential regulators of EEC differentiation (**Fig. 5F**). Consistent with this, key type I (e.g., ISG15) and type II IFN response genes (including both IRF1 and STAT1) were upregulated following lapatinib treatment (**Supplementary Fig 5H-I**).

**Figure 5.**
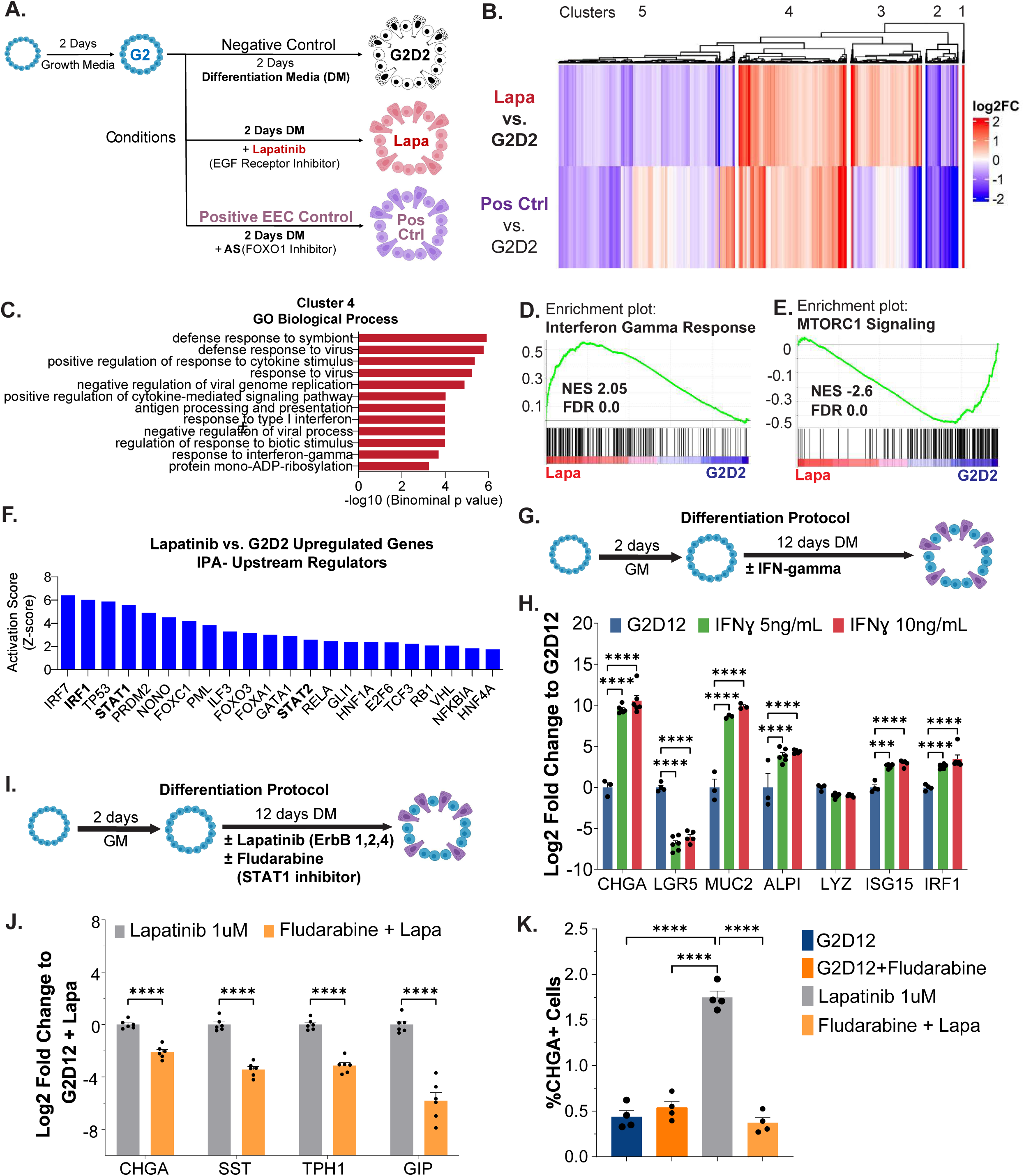
EGFR Inhibition Activates STAT1 Signaling to Drive EEC Differentiation in Human Organoids. (A) Schematic of experimental design for bulk RNA sequencing. Human duodenal organoids were grown for two days in growth media (GM) followed by 2 days in differentiation media (DM) alone (G2D2), with lapatinib (G2D2 + lapa), or with AS1842856 (positive control). (B) Heatmap of differentially expressed genes (DEGs) showing hierarchical clustering of genes shared between lapatinib- and AS-treated organoids compared to G2D2. Clusters 2 and 4 represent genes coordinately downregulated and upregulated, respectively, in both lapatinib- and AS-treated conditions relative to G2D2. (C) Gene Ontology (GO) biological process enrichment analysis of upregulated DEGs in Cluster 4, demonstrating enrichment of antiviral and inflammatory pathways, predominantly related to interferon (IFN) signaling. (D,E) Gene set enrichment analysis (GSEA) demonstrating enrichment of IFNγ response pathways among upregulated genes (D) and suppression of mTORC1 signaling, a known downstream target of EGFR, among downregulated genes (E) in lapatinib-treated organoids compared to G2D2. (F) Ingenuity Pathway Analysis (IPA) of upstream regulators of upregulated DEGs in lapatinib-treated organoids, identifying IRF1, STAT1, and STAT2 among the top predicted regulators. (G) Schematic of differentiation protocol for IFNγ treatment. Organoids were grown for two days in GM followed by 12 days in DM with or without increasing concentrations of IFNγ. (H) qPCR analysis of intestinal lineage and IFN response genes in organoids grown in G2D12 with increasing concentrations of IFNγ, normalized to *ACTB* and plotted as fold change relative to G2D12. CHGA = chromogranin A; LGR5 = leucine-rich repeat-containing G-protein coupled receptor 5; MUC2 = mucin 2; ALPI = intestinal alkaline phosphatase; LYZ = lysozyme; ISG15 = interferon-stimulated gene 15; IRF1 = interferon regulatory factor 1. Representative experiment showing n = 3–6 wells per condition from a single organoid line. (I) Schematic of differentiation protocol for combined lapatinib and fludarabine treatment. Organoids were grown for two days in GM followed by 12 days in DM with lapatinib with or without fludarabine (STAT1 inhibitor). (J) qPCR analysis of EEC markers in organoids treated with lapatinib alone or in combination with fludarabine, normalized to *ACTB* and plotted as log₂ fold change relative to G2D12 + lapatinib (Lapatinib). Representative experiment showing n = 6 wells per condition from a single organoid line. (K) Flow cytometry analysis of CHGA+ cells in organoids treated with G2D12, fludarabine alone, lapatinib alone, or lapatinib plus fludarabine. Representative experiment showing n = 4 wells per condition from a single organoid line. Bars show mean ± SEM; two-way ANOVA with Tukey correction for multiple comparisons (H, J); one-way ANOVA with Tukey correction for multiple comparisons (K). *p < 0.05; **p < 0.01; ***p < 0.001; ****p < 0.0001. Each experiment was repeated with at least three independent organoid lines.

Having identified IFN signaling as a candidate pathway, we next determined whether Type I or Type II IFN signaling was specifically required for EEC differentiation by exposing organoids to either RO8191 (an agonist for Type I IFN signaling) or IFN gamma (IFNγ) itself. Treatment with RO8191 confirmed activation of IFN target genes but failed to induce EEC markers, though a reduction in *LGR5* expression was observed (**Supplementary Fig 6A**). In contrast, IFNγ treatment led to a dose-dependent induction of EEC markers, with significant upregulation of *CHGA* and *MUC2* alongside confirmed activation of IFN target genes *ISG15* and *IRF1*, a reduction in the stem cell marker *LGR5*, and an increase in the enterocyte marker *ALPI* compared to G2D12 controls (**Fig 5G-H**). Consistent with these findings, IFNγ treatment also induced significant upregulation of EEC hormone genes including *MLN*, *SST*, and *GHRL* (**Supplementary Fig 6B**). Since IFNγ is well known to promote phosphorylation of STAT1, we next assessed lapatinib-treated human duodenal organoids by western blot analysis, which revealed induction of STAT1 phosphorylation within two hours (**Supplementary Fig 6C-D**)(*18*). To directly test whether STAT1 activation is functionally required for EGFRi-induced EEC differentiation, we co-treated organoid cultures with lapatinib and fludarabine, a small molecule inhibitor of STAT1 phosphorylation and nuclear translocation (**Fig 5I**). Fludarabine abrogated EGFRi-induced EEC differentiation, as evidenced by reduced expression of *CHGA*, *SST*, *TPH1*, and *GIP* and a marked reduction in CHGA+ cell numbers by flow cytometry (**Fig 5J-K).** Notably, *MUC2* expression was similarly reduced with co-treatment, supporting a broader role for the EGFRi-induced STAT1 signaling in secretory lineage differentiation (**Supplementary Fig 6E**). Collectively, these data position STAT1 as a critical mediator of EGFRi-induced EEC differentiation.

## DISCUSSION

Here, we demonstrate that EGFR inhibitors (among the most widely used targeted therapies in oncology), which frequently induce gastrointestinal toxicity, robustly drive ISC-to-EEC differentiation and hormone production through activation of STAT1 signaling. Using high-throughput screening, human intestinal organoids, and a patient cohort and murine model treated with erlotinib, we establish a direct mechanistic link between EGFR inhibition and EEC biology. These findings provide a mechanistic framework for EAD with direct implications for its clinical management and the development of novel mitigation strategies.

ErbB receptor signaling is a known regulator of ISC proliferation and lineage specification, with niche-derived EGF-family ligands influencing differentiation in crypts(*19*). Stromal ligands targeting specific ErbB receptors have been shown to promote specific phenotypes in human and murine studies(*19–22*). For example, an EGF/EREG (ErbB1/2) gradient along the crypt-villus axis seems to be important for ISC maintenance and specification of the absorptive lineage, while NRG1 (ErbB3/4) has been shown to facilitate crypt regeneration and a fetal-like repair program following injury(*20–22*). Our data further support a role for EGF signaling in ISC differentiation with inhibitors of multiple receptors (e.g., lapatinib targets ErbB1, 2 and 4 and erlotinib targets ErbB1) promoting EEC differentiation. Previous work in mouse organoids has shown that dampening EGFR or ERK signaling induces a reversible Lgr5+ ISC quiescence that is permissive for EEC differentiation; conversely, reinforced EGFR–ERK activity sustains ISC proliferation and promotes the absorptive lineage(*10*). Unlike previous reports relying on combinations of growth factor modulation or co-inhibition of parallel pathways, including WNT, our data reveal that the addition of EGFRi alone during human duodenal organoid differentiation promotes ISC differentiation and EEC hormone production. Together, these observations indicate a robust and direct regulatory role for ErbB family signaling in the modulation of EEC differentiation in human intestinal tissue.

IFN signaling has also been previously positioned as a modulator of ISC fate(*23–26*). In aged intestinal crypts, heightened IFNγ –STAT1 activity has been reported to increase differentiation of both absorptive and secretory lineages at the cost of the stem cell pool needed for regeneration(*24*). As a result, aging crypts exhibited increased expression of IFN-stimulated genes, including ISG15, also shown to be elevated in our studies. Interestingly, control of type I IFN signaling has been previously shown to be important in ISC fate decisions, as chronic IFN exposure, in the setting of deletion of the regulator IRF2, can bias ISC commitment toward the secretory lineage(*23*). Our organoids did not show transcriptional signs of differentiation in the setting of the Type I IFN activator RO8191, suggesting that a deeper analysis of this pathway, especially IRF2 expression and function, is warranted in future studies. Expanding on this, single cell multiomics and perturbation analyses in human ISC models engineered for endocrine differentiation (via NEUROG3 overexpression) have shown that early EEC commitment transiently induces IFN programs, with IRF1 emerging as a key marker of secretory progenitor cells(*12*). Building on this, we show that EGFRi in human duodenal organoids modulates STAT1 and induces type II IFN signaling targets, including the expression of IRF1, to promote secretory lineage and EEC differentiation.

Our data also suggest that there is increased expression of other differentiated lineage markers following EGFRi or IFNγ exposure, including goblet cells and enterocytes. These data are consistent with previous murine studies suggesting that activation of STAT1 through type II IFN signaling induces pan-differentiation in the intestine(*18, 27*). Interestingly, inhibition of STAT1 phosphorylation using fludarabine decreased the effect of EGFRi on EEC differentiation, but did not alter gene expression of goblet cell, Paneth cell, or enterocyte markers. This suggests that there are additional targets of EGFRi and IFNγ that function independently of STAT1 in the intestine to regulate differentiation.

EGFR inhibitors have been recognized as key regulators of inflammatory pathways outside of the intestinal epithelium and have been consistently linked to increased STAT1 activation through multiple established mechanisms(*28–31*). Resistance to EGFRi therapy in mutant lung cancer cell lines is associated with robust type I and type II IFN responses following treatment with these medications(*29*). In lung epithelial systems, EGFR inhibition promotes IFN signaling through multiple mechanisms, including DNA-sensing pathways and NF-κB signaling converging on TBK1–IRF3 activation, as well as modulation of NF-κB via miR-21 regulation (*28, 32*). In addition to IFN induction, EGFRi can enhance STAT1 phosphorylation through complementary mechanisms, including relief of EGFR–SHP2–mediated suppression(*33–35*). Consistent with this, we observe transcriptional upregulation of STAT1 after two days of lapatinib treatment, supporting engagement of a STAT1-dependent transcriptional program, in line with prior studies demonstrating that EGFR signaling regulates STAT1 and ISG expression in a context-dependent manner(*36, 37*). Together, these studies establish EGFRi as modulators of STAT1 signaling across systems. Consistent with this, our findings reproduce EGFRi-induced STAT1 activation in intestinal tissue and further demonstrate that this signaling axis is functionally required for EEC differentiation. While EGFRi promotes STAT1 activity, prior studies also indicate that, in the aging colon, IFN signaling can reciprocally regulate pathways downstream of EGFR signaling(*38*). In colonocyte monolayers, IFNγ preconditioning remodels EGFR signaling outputs, attenuating ERK phosphorylation and reprogramming ion transport(*38*). This bidirectional interplay highlights a broader EGFR–IFN–STAT1 signaling network with implications for epithelial differentiation and tissue homeostasis.

Although EAD is one of the most common side effects of EGFR inhibitors, how they induce intestinal toxicity is not fully understood(*16, 39*). For instance, EGFR inhibitors have been reported to acutely activate basolateral potassium channels and apical CFTR chloride channels in intestinal epithelium, increasing fluid secretion and stool water(*40*). Moreover, other reports emphasize mucosal atrophy and barrier alterations, including villus blunting with gefitinib or erlotinib in short-course mouse studies(*41–43*). Additional explanations have included immune activation (e.g., neratinib models responsive to budesonide), bile-acid malabsorption, and microbiome shifts—underscoring mechanistic heterogeneity by drug, dose, and schedule(*40, 44*). However, these explanations have not fully accounted for the high prevalence of EAD, its persistence following discontinuation of EGFRi, nor its resistance to standard antidiarrheal therapies(*39, 45*). Our findings implicate altered lineage specification—specifically, the expansion and activation of EEC differentiation—as a potential driver of EAD.

We show that ErbB inhibition enhances the production of pro-motility hormones (e.g., 5HT and MLN), potentially inducing functional changes in GI motility without overt epithelial damage. In parallel, our animal models exhibited reduced food and water intake, which may reflect increased levels of satiety-associated hormones such as GLP-1 and PYY, thereby contributing to weight loss observed in patients. Inhibition of STAT1 signaling, through JAK1/2 blockade, has also been shown to ameliorate EGFRi–associated toxicities such as alopecia, where loss of EGFR signaling induces a hypersensitive JAK–STAT1 inflammatory program, highlighting STAT1 as a potential therapeutic target that may also be relevant for EAD(*37*). Notably, topical JAK inhibition has recently demonstrated clinical efficacy in ameliorating EGFRi-induced skin rash in both rodent models and cancer patients38, further highlighting STAT1 as a convergent therapeutic target across multiple EGFRi-associated toxicities that may also be relevant for EAD(*31*). These insights not only contribute to the potential mechanisms underlying EAD but open new avenues for targeted mitigation strategies, including manipulation of inflammatory pathways such as type II IFN signaling.

Despite these advances, our study has limitations. Although our human organoid models provide important translational insights, they lack the natural gradients of key growth factors found *in vivo*. Nevertheless, in our model, we demonstrate that EEC differentiation can occur even in the presence of WNT and other essential signals, making our system closely representative of homeostatic physiology. Moreover, while we link EEC differentiation and hormonal secretion to diarrhea in murine models, other mechanisms may also contribute to the observed phenotypes. Conditional knockout models lacking EECs would be ideal for determining the dependence of EAD on EEC physiology; however, these models also exhibit baseline phenotypes such as weight loss and malabsorption(*46*), which could confound interpretation. Additionally, although our patient cohort demonstrated an association between EGFRi use and increased EEC hormone levels, we cannot fully account for confounding factors (such as concomitant use of medications and/or dietary factors) that may influence hormone levels. As the patient samples were collected during routine clinical care, it was not possible to directly correlate hormone levels with the incidence of diarrhea, nor were direct measurements of EEC numbers in intestinal tissue available. A prospective study comparing serum hormone levels before and after EGFRi therapy in a controlled setting would more effectively address these limitations.

## MATERIALS AND METHODS

### 1. Organoid Culture

#### 1.1. Tissue Sources and Lines Used

Human intestinal organoid lines were generated from de-identified biopsies obtained from patients undergoing endoscopy at Boston Children’s Hospital. Samples were taken from tissue regions without gross pathology. Consent from guardians and assent from donors were collected in accordance with IRB approval (Protocol number IRB-P00000529). Organoid lines were derived from adolescent donors aged 13-19 years (n=9; 5 male, 4 female). Donor age and sex were unknown to researchers when experiments were being performed. All experiments used a minimum of three independent organoid lines.

#### 1.2. Crypt Isolation and Organoid Establishment

Biopsies were either processed immediately in DMEM/F12 or cryopreserved in 10% FBS with DMSO. To generate organoids, intestinal crypts were isolated from fresh or thawed samples by rinsing in DMEM/F12 and digesting in 2 mg/mL Collagenase Type I prepared in Hank’s Balanced Salt Solution. Tissue fragments were agitated with a motorized pestle, then incubated at 37 °C for 40 minutes. At the midpoint, digestion was aided by pipetting with a bent pipette tip. Following incubation, samples were centrifuged at 800 × g for 5 minutes and supernatants were aspirated. Crypts were resuspended in 150 μL Matrigel, with 40 μL seeded into three wells of a 24-well plate and polymerized at 37 °C for 10 minutes before culture medium was added.

#### 1.3. Culture Conditions and Media

Established duodenal organoids were expanded in growth medium (GM) composed of 65% L-WRN conditioned medium, 30% DMEM/F12, and supplements including Glutamax (1%), N-2 (1%), B-27 (1%), 10 mM HEPES, Primocin (100 μg/mL), Normocin (100 μg/mL), A83-01 (500 nM), N-acetyl-cysteine (500 μM), recombinant murine EGF (50 ng/mL), [Leu15] Gastrin I (50 nM), nicotinamide (10 mM), and SB202190 (10 μM). Media was refreshed every two days, and organoids were passaged every 6–8 days.

For differentiation, organoids were cultured in GM for 2 days post-passage to allow for stem cell expansion before being switched to differentiation medium (DM). DM contained 50% L-WRN conditioned medium, 45% DMEM/F12, and the same supplements as GM (except for SB202190 and nicotinamide), with the addition of DAPT (20 μM), betacellulin (20 ng/mL), PF06260933 (6 μM), tranylcypromine (1.5 μM), and Tubastatin-A (10 μM; removed after 48 hours). Media was replaced every two days. Organoids were harvested for downstream assays after 4–14 days of culture.

#### 1.4. Passaging and Maintenance

For passage, Matrigel domes were detached by adding 500 μL of 5 mM EDTA/PBS to each well. The suspension was transferred to a 15 mL conical tube and incubated on ice for 1 hour with inversion every 20 minutes. Organoids were pelleted at 500 × g for 5 minutes at 4 °C, the supernatant was removed, and organoids were fragmented mechanically using a bent pipette tip. Cells were resuspended in a 1:2 dilution of Matrigel and replated as 40 μL domes in 24-well plates. Domes were polymerized at 37 °C before adding 500 μL GM.

#### 1.5. Small Molecule Inhibitor Experiments

For inhibitor studies, organoids were expanded in GM for 2 days post-passage, then transitioned to DM supplemented with small molecules as described in **Supplementary Table 3**. Media was replaced every 48 hours, and organoids were collected after 4–14 days for analysis.

### 2. High-Throughput Screening

#### 2.1. Cell Culture Conditions

Organoid lines were expanded from previously characterized lines and screened for their ability to differentiate into EECs by immunofluorescent staining. Lines younger than passage 20 were selected to minimize variability in differentiation potential.

For plating, organoids were released from Matrigel using 5 mM EDTA/PBS and incubated on ice for 40–60 minutes. Organoids were pelleted at 500 × g for 5 minutes at 4 °C, with additional centrifugation performed as needed. Pellets were resuspended in 1200 μL Matrigel and fragmented by pipetting with a bent 1000 μL tip for 1–2 minutes. The suspension was combined with 4000 μL Matrigel and 250 μL GM, yielding 5450 μL total, sufficient to seed eight 96-well plates.

5 μL of the suspension was dispensed into each well of the 96-well plates using a multichannel repeater pipette, ensuring domes were formed without Matrigel contacting the well walls. Plates were incubated at 37 °C for 10 minutes to allow for polymerization, then 150 μL GM was added. After two days, GM was replaced with DM supplemented with library compounds.

#### 2.3. Compound Library and Screening Protocol

High-throughput screening was performed using the MedChemExpress small molecule library, pre-formatted into five 384-well Echo (Beckman Coulter)-compatible LP-0200 plates. Each plate was subdivided into quadrants and transferred into twelve 96-well polypropylene plates, generating three replicate plates per quadrant. Plates were heat-sealed and stored at -20 °C until used.

To perform the screen, each 96-well organoid plate included a negative control column (DM only) and a positive control column (DM + AS). To begin the study, one set of library plates was thawed for 15 minutes at room temperature under desiccated conditions, centrifuged at 1000 rpm, and unsealed. Compounds were diluted by adding 120 μL fresh medium to each well using a Bravo liquid handler (Agilent), followed by mixing. Medium was removed from assay plates containing organoids using a Multidrop Combi rapid plate dispenser (Thermo Fisher), and 100 μL of fresh medium was added. From each well of the compound plate, 50 μL of compound-containing medium was dispensed into the assay plates for a final concentration of 5 μM. Plates were handled only by their edges to avoid static-induced redistribution of small volumes. The screening schedule was as follows: day 1, organoid plating; day 3, first compound addition; day 6, medium exchange and second compound addition; day 9, medium exchange and third compound addition; and day 11, fixation and staining as described below. Plates were maintained at 37 °C, 5% CO₂ throughout the assay.

#### 2.2. Staining and immunofluorescence

On day 11, organoids in Matrigel were fixed with 4% paraformaldehyde (150 μL/well) for 30 minutes at 37 °C with shaking (70–80 rpm). Samples were permeabilized with 0.5% Triton X-100 in TBST for 30 minutes at room temperature, then blocked with 10% FBS/1% BSA in TBST for 30 minutes. Organoids were incubated overnight at 4 °C with primary antibody against CHGA (mouse, Dako, 1:100) prepared in 0.25% Triton X-100/10% FBS/1% BSA/TBST. The following day, samples were washed three times in 0.2% Triton X-100/TBST for 10 minutes each, then incubated with secondary antibody (donkey anti-mouse Alexa Fluor 568, 1:400) and DAPI (1:1000) for 2 hours at room temperature in the dark. After three additional washes, organoids were resuspended in 100 μL PBS for imaging. Liquid handling was performed using a Multidrop Combi rapid plate dispenser (Thermo Fisher).

#### 2.4. Hit Identification and Selection Criteria

Imaging was performed using the ImageXpress Micro Confocal Laser (Thermo Fisher). Fluorescence channels (DAPI and 568 nm) were first normalized to positive and negative control wells to establish consistent exposure parameters for both fluorophores across plates. Based on these parameters, EEC differentiation was quantified using CHGA immunofluorescence (measured at 568 nm). A compound was considered to promote differentiation if it exceeded the negative control in both of the following metrics: 1) the proportion of CHGA-positive organoids relative to total organoids, and 2) the proportion of CHGA-positive signal relative to total DAPI-positive area. The first parameter assessed consistency between organoids, while the second controlled for differences in total cell number. A minimum threshold of 0.01 was applied to classify CHGA positivity. CHGA signal was quantified as feature counts and normalized to total organoid area to account for differences in organoid size and cell number. The percentage of CHGA-positive cells per well was calculated as: [%CHGA+ Cells/DAPI Area = (CHGA feature count / total organoid area) × 100]. Compounds were then ranked based on this normalized CHGA signal (%CHGA+ Cells/DAPI Area), and distributions were visualized across all screened conditions.

### 3. Quantitative PCR

Total RNA was isolated from individual wells using TRI® Reagent and the Direct-zol RNA kit according to the manufacturer’s instructions. RNA concentration was determined with a NanoDrop 1000 spectrophotometer (Life Technologies). Reverse transcription was performed using the High-Capacity cDNA Reverse Transcription Kit. qPCR reactions were carried out on a QuantStudio 6 Flex thermocycler (Thermo Fisher) using TaqMan probes listed in **Supplementary Table 4**. 18S or ACTB were used as housekeeping genes, and relative expression was calculated by the 2−ΔCt method with a cutoff Ct of 40.

### 4. Flow Cytometry

Organoids were released from Matrigel using 5 mM EDTA/PBS on ice for 10 minutes, pelleted at 800 × g for 5 minutes at 4 °C, and dissociated in 500 μL TrypLE Express for 10 minutes at 37 °C. Single-cell suspensions were obtained by pipetting with a bent tip. Cells were diluted in 20% FBS/DMEM-F12 and centrifuged. Dead cells were labeled with DAPI (1:1000) in 2% FBS/2 mM EDTA/DMEM for 10 minutes at room temperature. Cells were then fixed in 1% PFA, washed, and permeabilized in 0.2% Tween-20/2% FBS/PBS for 15 minutes at 37 °C. Antibody staining was performed for 30 minutes on ice using reagents in **Supplementary Table 5**. Samples were resuspended, filtered through a 37 μm mesh, and analyzed on a BD LSRFortessa with BD FACSDiva and FlowJo software.

### 5. Immunofluorescence

Organoids were collected as above, centrifuged at 500 × g for 5 minutes, and fixed in 4% PFA for 20 minutes on ice with agitation. After centrifugation, organoids were washed and stored in PBS at 4 °C until staining (≤7 days). Organoids were permeabilized in 0.3% Triton X-100 in PBS for 20 minutes at room temperature, then blocked in 5% BSA for 1.5 hours. Primary antibodies described in **Supplementary Table 5** were diluted in 5% BSA/0.1% Triton X-100 and incubated overnight at 4 °C. Organoids were washed five times in 0.1% Triton X-100, incubated with secondary antibodies for 2 hours at room temperature, washed again, and mounted in ProLong Gold Antifade. Slides were sealed, dried overnight, and stored at 4 °C.

Organoid immunofluorescence quantification was conducted as previously described(*14*). Briefly, organoids were imaged using an EVOS FL 2 Auto microscope (Thermo Fisher) at 2× magnification, and stitched images were processed in Fiji. DAPI channels were converted to 8-bit grayscale, smoothed with a Gaussian blur (radius = 4), and manually thresholded to separate individual organoids. Segmentation was refined using Watershed and Find Edges functions, and organoids were quantified with *Analyze Particles* (size = 4000–100,000; circularity = 0.30–1.00). Outlines from DAPI masks were applied to corresponding fluorescent channels, which were thresholded manually to define positive staining. Mean fluorescence intensity was extracted from each ROI, and organoids with nonzero mean values were classified as positive.

### 6. Hormone Quantification (ELISA)

Organoids were plated in 48-well plates (25 μL Matrigel/well) and differentiated as above with the addition of erlotinib or lapatinib. To stimulate secretion, forskolin was added 24 hours before conditioned media was collected, flash frozen, and stored at −80 °C. All conditions included Diprotin A (100 μM) to prevent hormone degradation. Quantification was performed using ELISA kits for serotonin, GIP, Motilin, Somatostatin, PYY, and GLP-1 (**Supplementary Table 5**). Samples were normalized to DNA content, measured after NaOH lysis and Tris neutralization, followed by quantification with a NanoDrop 1000 spectrophotometer.

For human studies, we used de-identified serum samples from a cohort of 94 patients (Supplementary Fig. 2). Samples were previously collected prospectively under IRB protocols #19-201 and #02-180 (“Collection of Specimens and Clinical Data for Subjects with Proven or Suspected Thoracic Malignancies”) at Dana-Farber Cancer Institute and stored at −80 °C in the Christiani Lab. Prior to analysis, frozen serum samples were thawed and processed using human ELISA kits to quantify 5HT, MLN and SST (**Supplementary Table 5**).

For mouse studies, blood was collected via cardiac puncture into protease inhibitor lined syringes and placed in EDTA tubes for plasma or Eppendorf tubes for serum. Samples were spun down at 1000 x g for 10 minutes at 4°C (plasma) or room temp after clotting (serum). To quantify levels of PYY and total GLP-1 in plasma, custom Mesoscale U-plex plates were utilized and run in duplicate as per manufacturer’s instructions. To quantify 5HT, SST, GHRL and GIP, serum samples were run on mouse 5HT, human GIP or human SST ELISA plates as per manufacturer’s instructions.

### 7. Western Blotting

#### 7.1. Organoid Collection

Organoids were released from Matrigel with 5 mM EDTA/PBS on ice for 10 minutes. Organoids from 3–4 wells were pooled into 15 mL conical tubes, pelleted by centrifugation at 800 × g for 5 minutes, and rinsed with PBS. Samples were transferred into 1.5 mL tubes, centrifuged at maximum speed for 5 minutes, and excess liquid was carefully removed. Pellets were flash-frozen in liquid nitrogen and stored at –80 °C.

#### 7.2. Protein Extraction

Pellets were thawed on ice and lysed in RIPA buffer containing protease and phosphatase inhibitors. Lysates were briefly sonicated (10 seconds, 20% amplitude) and kept on ice throughout processing. Protein concentration was measured using the Pierce BCA Protein Assay Kit.

#### 7.3. SDS-PAGE and Transfer

Equal protein quantities were mixed with a loading buffer, heated at 95 °C for 5 minutes, and separated on SDS–polyacrylamide gels. Electrophoresis was carried out at 80 V for 20 minutes followed by 100 V until the dye front reached the bottom. Proteins were transferred to PVDF membranes using the Trans-Blot Turbo Transfer System (Bio-Rad).

#### 7.4. Antibody Incubation

Membranes were blocked for 1–2 hours in 1% BSA in PBS with sodium azide. Membranes were cut by molecular weight where necessary to allow probing with multiple antibodies. Primary antibodies described in **Supplementary Table 5** were diluted in 1% BSA/PBS and incubated overnight at 4 °C. After three 5-minute washes in TBST, membranes were incubated with HRP-conjugated secondary antibodies in TBST for 1.5–3 hours at room temperature.

#### 7.5. Detection and Quantification

Signal was detected using enhanced chemiluminescence (ECL) reagents, imaged with a ChemiDoc system. Exposure times were adjusted as needed (maximum 5–8 minutes). Protein levels were quantified using ImageJ and normalized to housekeeping proteins (β-actin, vinculin).

### 8. In Vivo Animal Studies

#### 8.1. Animal Models and Husbandry

Male and female C57BL/6J mice were purchased from Jackson Laboratories and housed in a specific-pathogen-free (SPF) facility at Boston Children’s Hospital with a 12:12 light-dark cycle and *ad libitum* food and water. Animals were used at 8-12 weeks age. Both sexes were included in all experiments, and no significant sex differences were observed in the measured outcomes. All studies were approved by the Institutional Animal Care and Use committee (IACUC) at Boston Children’s Hospital (Protocol #00002432 and #00001721).

#### 8.2. Treatment Regimens

Animals were treated once daily by oral gavage with either vehicle (40% Captisol® [sulfobutylether-β-cyclodextrin, SBE-β-CD]) or erlotinib (120 mg/kg).(*17*) Erlotinib was formulated in 40% Captisol®, and dosing solutions were freshly prepared daily prior to administration.(*34*)

#### 8.3. Measurement of Gastrointestinal Motility

1-hour fecal pellet output and stool water were assessed as previously described (Rao et al., 2017; Shepherd et al., 2024). Briefly, animals were placed in empty cages at 9AM for 1 hour. Pellets were immediately removed after passage and placed in pre-weighed Eppendorf tubes. After one hour, animals were returned to their home cage and the tube containing wet stool weighed. The fecal pellets were then dried at 55°C for 72 hours and the tube weighed again to determine the percentage of water in the fecal contents.

#### 8.4 Measurement of Food and water consumption

Food and water intake was tracked within home cages. Each cage was provided with a 100 mL feeder bottle (Braintree Scientific, MA, WTRBTL S-BL) and a custom-made food metal hopper with a tray to catch chow remnants at the start of the experiment. Amount of food (in grams) and water (in mL) consumed was assessed every morning, along with animal body weight, and normalized by number of mice per cage to get the average food/water consumption of a mouse in each cage.

#### 8.4 Sample Collection and Processing

For EEC quantification, animals were euthanized via CO2 inhalation, and the intestine was removed and flushed with ice-cold PBS. 0.5 cm pieces of tissue were added to Trizol and processed for RNA using TRI® Reagent and the Direct-zol RNA kit according to the manufacturer’s instructions. Remaining tissue was then Swiss rolled. Tissue was fixed (overnight in 10% neutral buffered formalin) and paraffin embedded for downstream analysis.

Immunofluorescence measurement of CHGA-positive cells was done as previously described(*47*); antibodies used are displayed on **Supplementary Table 5**. For whole tissue CHGA quantification, five images were acquired from each intestine per animal using a Nikon upright Eclipse 90i microscope. For each image, three Z-stacks were collected and deconvoluted using NIS-Elements Nikon software to obtain optimal resolution.

Intestinal images were stitched together and adjusted for brightness and contrast using ImageJ software. Brightness levels were optimized to enhance visibility without causing pixel overexposure, and regions containing paraffin folding or nonspecific background were removed using ImageJ.

Images were exported in PNG format (separate files for each channel) and imported into Photoshop (version 25.2.0) as individual layers. Quantification was performed using one complete intestinal section per mouse. Epithelial tissue was delineated using an anti–E-cadherin antibody (DECMA-1, Abcam, ab11512), and CHGA–positive cells were detected using a rabbit polyclonal anti–Chromogranin A antibody (Proteintech, 10529-1-AP).

Quantification was based on pixel area using the histogram function. Specifically, the pixel area corresponding to CHGA–positive regions within the E-cadherin–defined epithelial compartment was measured and normalized to the pixel area of DAPI staining to account for variations in cell number. This normalization ensured accurate comparison across samples.

### 9. Bulk RNA Sequencing

Bulk RNA sequencing was performed on organoids cultured as described above, with samples collected at the indicated experimental time points under growth conditions, differentiation media (DM), and DM supplemented with either lapatinib (1.0 µM) or AS1842856 (100 nM). Total RNA was isolated using a column-based purification method according to the manufacturer’s instructions. RNA quantity and purity were assessed by spectrophotometry, and RNA integrity was evaluated by capillary electrophoresis prior to library preparation. Complementary DNA (cDNA) libraries were generated from poly(A)-enriched RNA using a stranded library preparation workflow, followed by adapter ligation, amplification, and quality assessment of library size distribution and concentration before sequencing on an Illumina platform to generate paired-end reads. Raw sequencing reads underwent quality control assessment and adapter trimming prior to alignment to the human reference genome using STAR, and gene-level counts were generated using featureCounts(*48, 49*). Differential gene expression analysis between conditions was performed using DESeq2(*50*). All genes were visualized in volcano plots using the EnhancedVolcano R package. Log2 fold-changes of the common differentially expressed genes (DEGs) between the two groups were displayed in a heatmap generated with pheatmap. Downstream transcriptomic analyses included pathway and functional enrichment analyses using Ingenuity Pathway Analysis (IPA), Gene Ontology (GO) Biological Process enrichment, Kyoto Encyclopedia of Genes and Genomes (KEGG) pathway analysis, and Gene Set Enrichment Analysis (GSEA)(*51*). GO and KEGG enrichment tests were performed using clusterProfiler v4.6.2(*52*). GO terms or pathways were considered significantly enriched when the adjusted p-values with Benjamini-Hochberg correction was smaller than 0.05. Bar plots of significant pathway or GO terms were generated using the enrichplot R package. GO terms were summarized in treemaps to reduce redundancy among semantically similar terms using rrvigo(*53*). Bulk RNA-seq data have been deposited in the Gene Expression Omnibus (GEO) under accession number GSE329493.

### 10. Statistical Analysis

All experiments were repeated using at least three different human organoid lines, with representative data from a single line shown, unless otherwise noted. For qPCR, flow cytometry, immunofluorescence, Western blot and ELISA, each condition was performed using pooled enteroids from 1-5 wells, unless otherwise noted, with each pooled group acting as a single sample. Prior to statistical analysis, all qPCR data were transformed using log2. When analyzing only two conditions, statistical significance was determined by unpaired, two-tailed Student’s t test. When analyzing more than two conditions, statistical significance was determined by either one-way or two-way ANOVA, followed by Tukey post hoc analysis. Unpaired t-tests were used to compare column analysis from flow cytometry, ELISA and western blot experiments. Welch’s t-test was used for selected mouse experiments to account for unequal variance between treatment groups and to provide a more robust comparison for datasets with relatively small sample sizes.

## Acknowledgment

We thank members of the Breault laboratory and the Division of Endocrinology at Boston Children’s Hospital for helpful comments and suggestions. We appreciate the support from the HDDC Histology and Organoid Cores, the BCH Bioinformatics Core, and the ICCB-Longwood Screening Facility at Harvard Medical School. This work was supported by funding from the Adolph Coors Foundation (to DTB) and the American Gastroenterology Association Pilot Research Award (AGA2024-21-1) (to GPR), and is the result of NIH funding R01DK135707 (to MR), K08DK134885 (to DZ), and P30DK034854 (to DTB), in whole or in part, and is subject to the NIH Public Access Policy. Through acceptance of this federal funding, the NIH has been given a right to make the work publicly available in PubMed Central. Images were made with BioRender.

## Abbreviations

CCK: Cholecystokinin
CHGA: Chromogranin A
EAD: EGFR inhibitor-associated diarrhea
EGF: Epidermal Growth Factor
EGFR: Epidermal Growth Factor Receptor
EGFRi: Epidermal Growth Factor Receptor inhibitor
EEC: Enteroendocrine Cell
GHRL: Ghrelin
GI: Gastrointestinal
GIP: Gastric Inhibitory Peptide
GLP-1: Glucagon-Like Peptide 1
IFN: Interferon
IRF1: Interferon Regulatory Factor 1
ISC: Intestinal Stem Cell
JNK: c-Jun N-terminal Kinase
LGR5: Leucine-rich repeat-containing G-protein coupled receptor 5
MLN: Motilin
PYY: Peptide YY
SST: Somatostatin
STAT1: Signal Transducer and Activator of Transcription 1
T2DM: Type II Diabetes Mellitus
TPH1: Tryptophan Hydroxylase 1
WNT: Wingless-Related Integration Site

## SUPPLEMENTARY MATERIALS

**Supplementary Figure 1.**
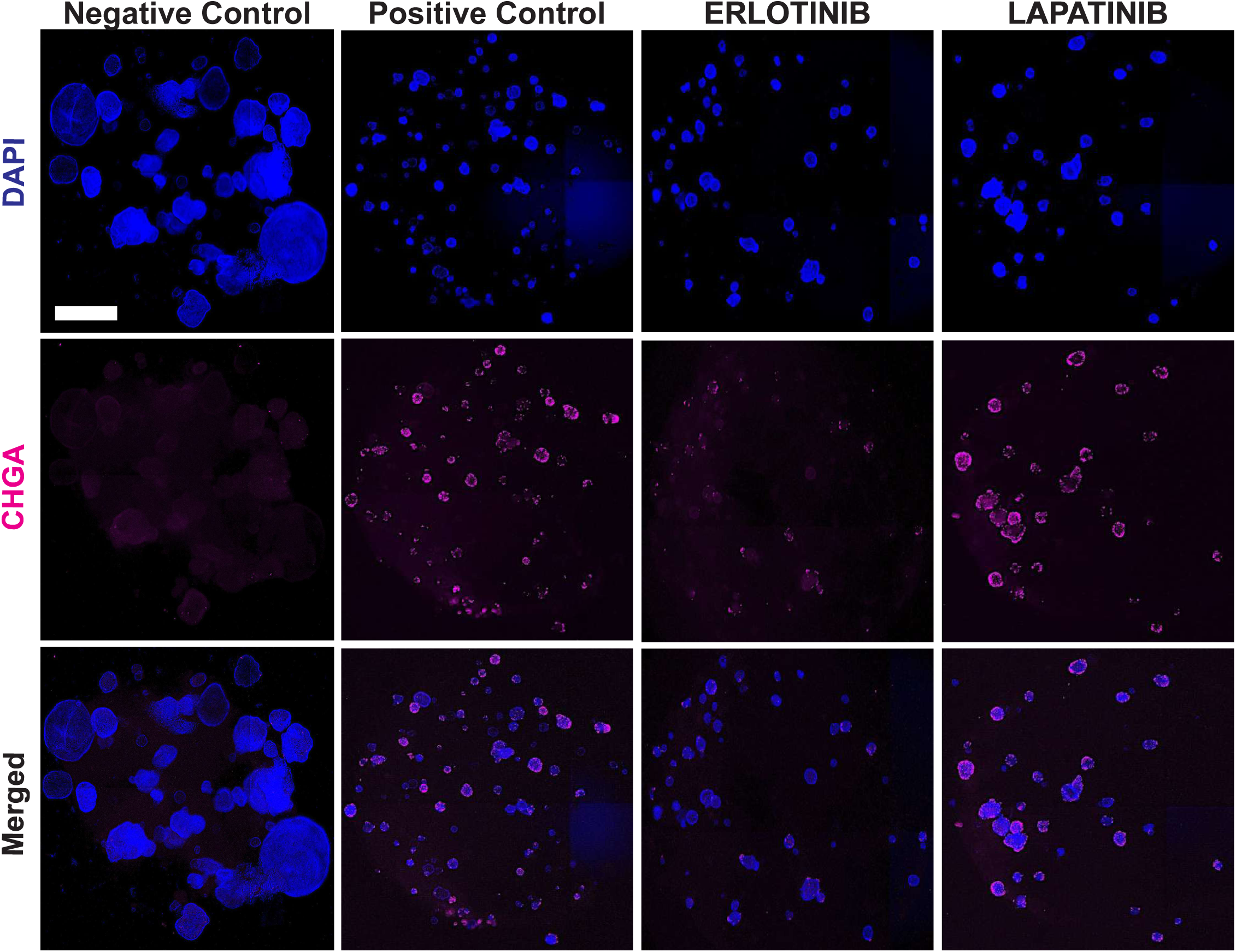
EGFR Inhibitors Induce CHGA Expression Comparable to Positive Control in Human Organoids. Representative immunofluorescence staining of human duodenal organoids following differentiation in base differentiation media (negative control), AS1842856 (positive control), erlotinib, or lapatinib. CHGA (magenta) and nuclei (DAPI, blue) are shown, along with merged images. Organoids were grown for two days in growth media (GM) followed by nine days in differentiation media (DM) with indicated treatments. Scale bar = 1 mm.

**Supplementary Figure 2.**
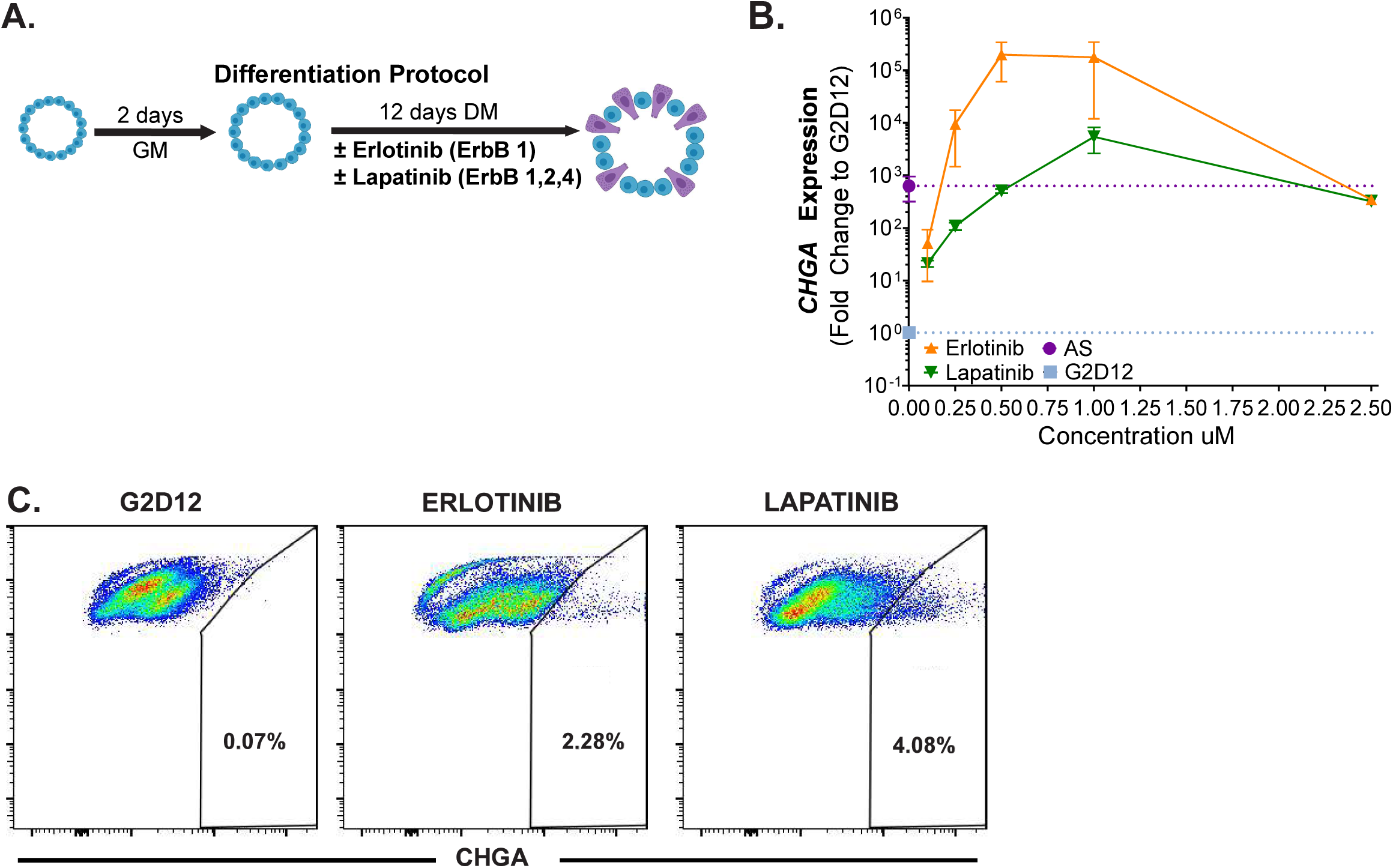
Dose-Dependent Induction of CHGA Expression by EGFR Inhibitors in Human Organoids. (A) Schematic of our differentiation protocol. Human duodenal organoids were grown for two days in growth media (GM) followed by 12 days in differentiation media (DM) with and without erlotinib or lapatinib. (B) qPCR analysis of *CHGA* expression across increasing concentrations of erlotinib and lapatinib, normalized to *ACTB* and plotted as fold change relative to G2D12. AS1842856-induced *CHGA* expression is shown as a positive control. (C) Representative flow cytometry plots of CHGA⁺ cells in organoids grown in G2D12, G2D12 with erlotinib, and G2D12 with lapatinib. Percentages of CHGA⁺ cells are indicated.

**Supplementary Figure 3.**
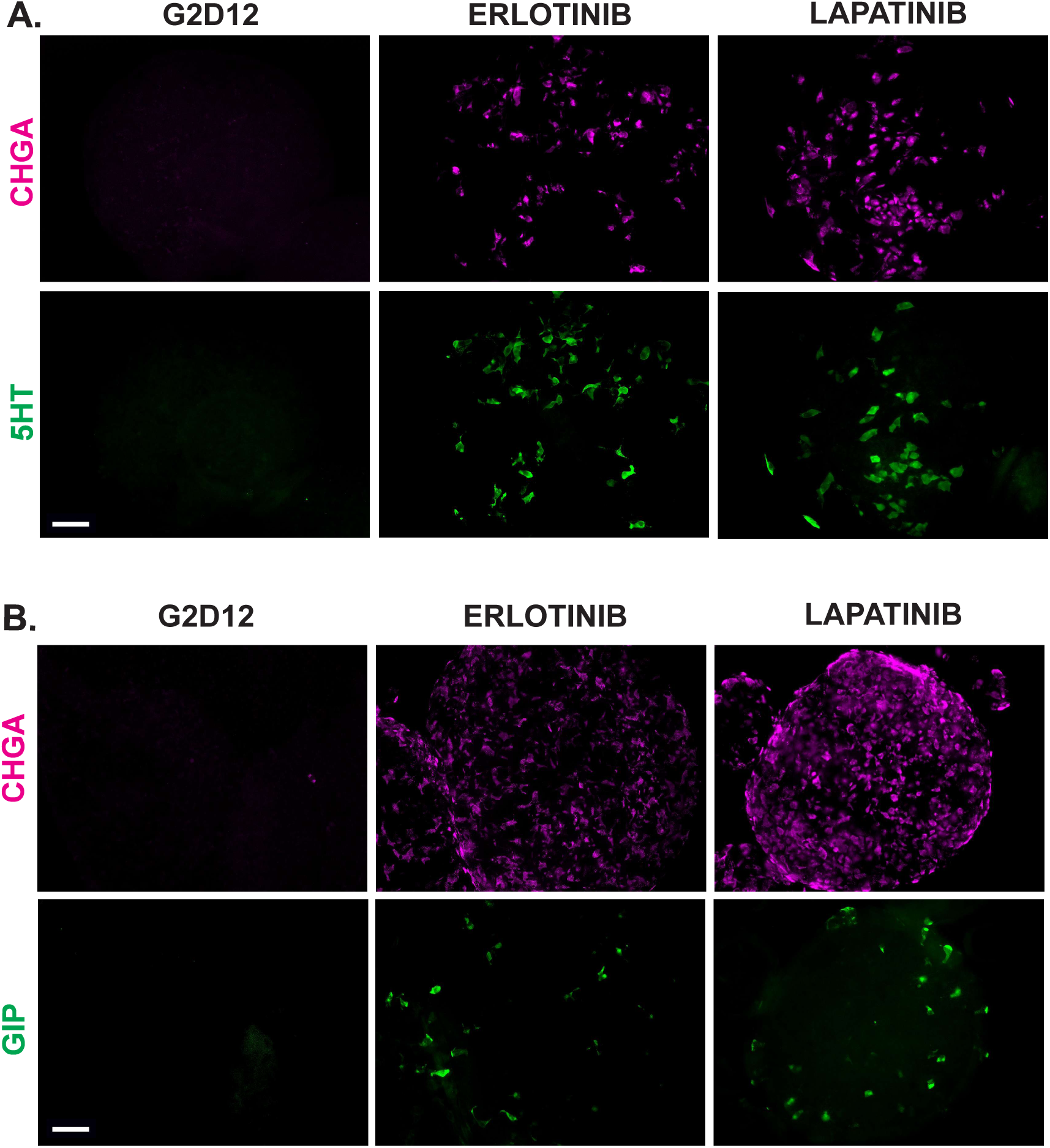
EGFR Inhibitors Induce 5HT- and GIP-Producing EECs in Human Organoids. (A,B) Representative immunofluorescence staining of human duodenal organoids grown in G2D12, G2D12 with erlotinib, and G2D12 with lapatinib. Individual channels are shown for CHGA (magenta) with 5HT (green, (A)) and GIP (green, (B)), corresponding to merged images shown in Figure 2C,D. Scale bars = 50 µm.

**Supplementary Figure 4.**
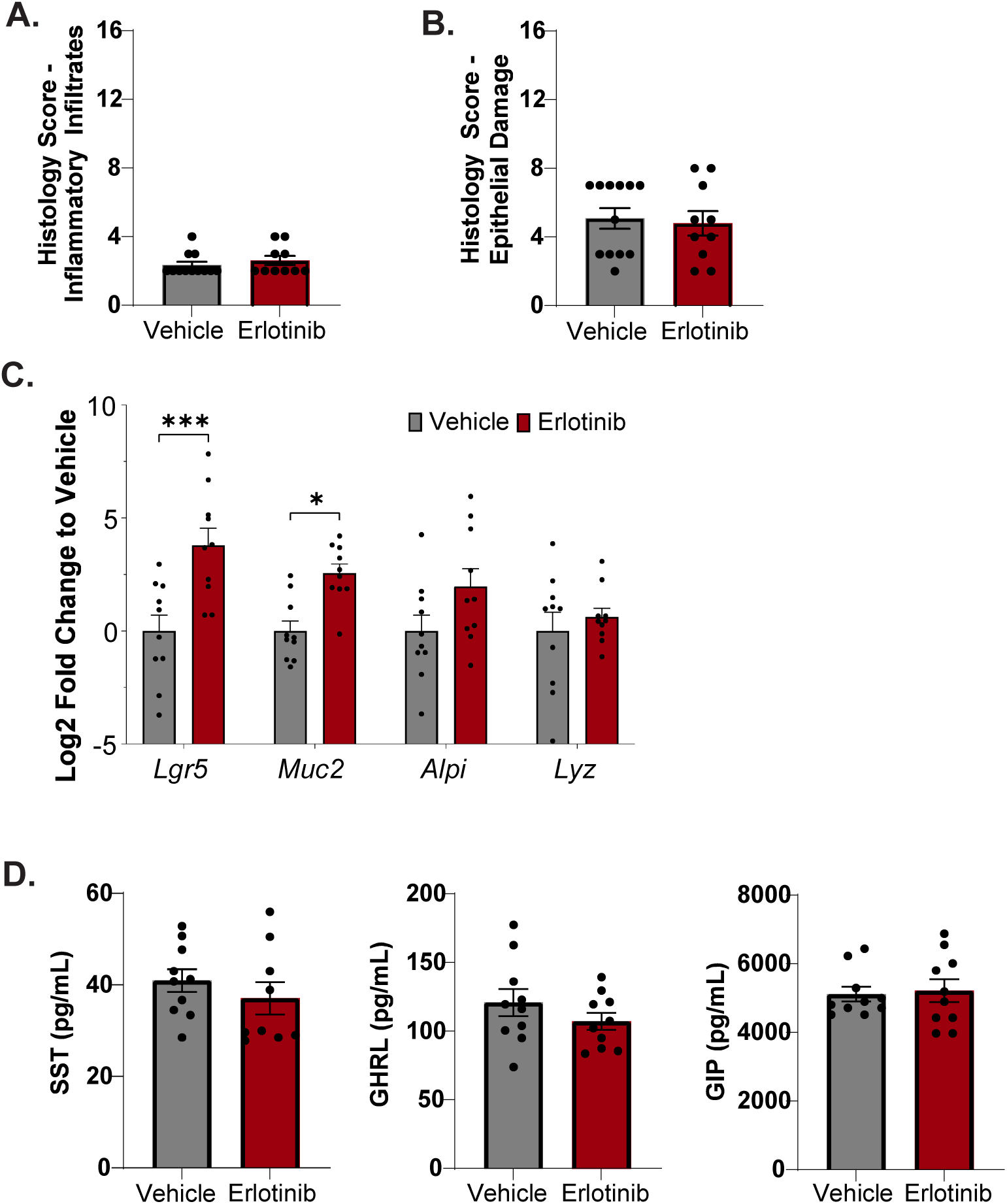
Additional Histological, Gene Expression, and Hormone Analyses in Erlotinib-Treated Mice. (A,B) Histological scoring of inflammatory infiltrate (A) and epithelial damage (B) in duodenum from vehicle-and erlotinib-treated mice (n = 10 mice per group). (C) qPCR analysis of additional intestinal lineage markers in duodenal tissue from vehicle- and erlotinib-treated mice, normalized to *Actb* and plotted as log₂ fold change relative to vehicle-treated mice. Lgr5 = leucine-rich repeat-containing G-protein coupled receptor 5; Muc2 = mucin 2; Alpi = intestinal alkaline phosphatase; Lyz = lysozyme. Representative experiment from one cohort of mice. (D) Serum levels of SST, GHRL, and GIP measured by ELISA from serum collected on day 7. Representative experiment from one cohort of mice. Bars show mean ± SEM; Welch’s t-test (A, B, D); two-way ANOVA with Tukey correction for multiple comparisons (C). *p < 0.05; ***p < 0.001. Experiments were repeated in three independent cohorts of mice.

**Supplementary Figure 5.**
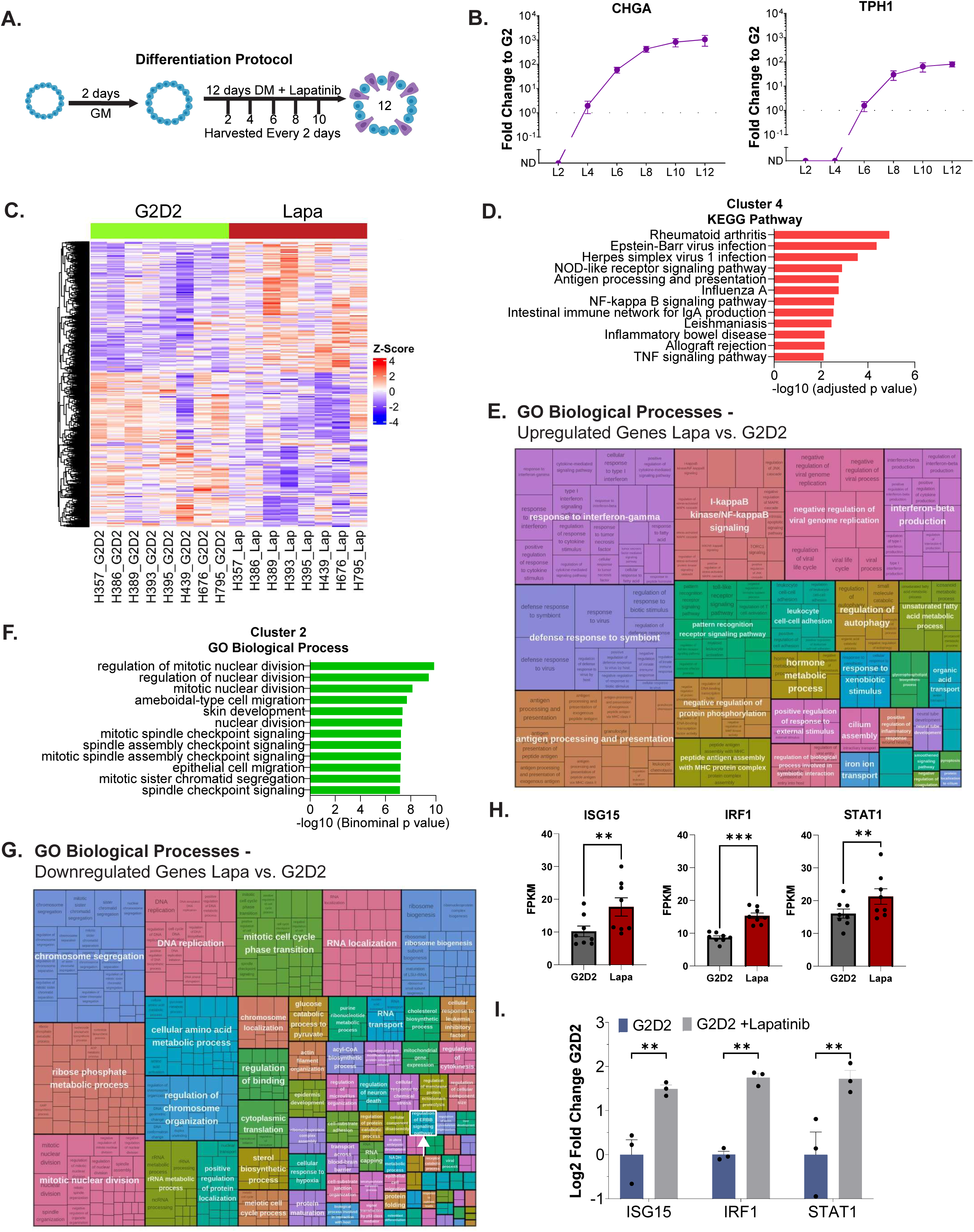
Temporal and Transcriptomic Analyses of EGFR Inhibition in Human Organoids. (A) Schematic of differentiation protocol for time-course analysis. Human duodenal organoids were grown for two days in growth media (GM) followed by 12 days in differentiation media (DM) with lapatinib, with samples collected every 2 days. (B) qPCR analysis of *CHGA* and *TPH1* expression over time in 1uM lapatinib-treated organoids, normalized to *ACTB* and plotted as fold change relative to G2 (dotted line). Representative experiment showing n = 4 wells per condition from a single organoid line. (C) Heatmap of differentially expressed genes (DEGs) in organoids grown for 2 days in GM followed by 2 days in DM (G2D2) with or without lapatinib, showing hierarchical clustering across independent organoid lines. (D) KEGG pathway enrichment analysis of upregulated genes in Cluster 4, highlighting pathways related to immune and antiviral responses. (E) Gene Ontology (GO) biological process enrichment of upregulated DEGs in lapatinib-treated organoids compared to G2D2, demonstrating enrichment of IFN-related and antiviral pathways. (F) GO biological process enrichment of downregulated genes in Cluster 2, highlighting pathways associated with cell cycle and mitotic processes. (G) GO biological process enrichment of downregulated DEGs in lapatinib-treated organoids compared to G2D2, including pathways associated with EGFR/ERBB signaling (white arrow) and related downstream processes. (H) Expression of representative IFN response genes (ISG15, IRF1, STAT1) in G2D2 and lapatinib-treated organoids from RNA-seq study. Data represent averages from eight different organoid lines. (I) qPCR validation of IFN response genes, normalized to *ACTB* and plotted as log₂ fold change relative to G2D2. Representative experiment showing n = 3 wells per condition from a single organoid line. Bars show mean ± SEM; unpaired, two-tailed Student’s t-test (H); two-way ANOVA with Tukey correction for multiple comparisons (I). **p < 0.01; ***p < 0.001. Each experiment was repeated with at least three independent organoid lines.

**Supplementary Figure 6.**
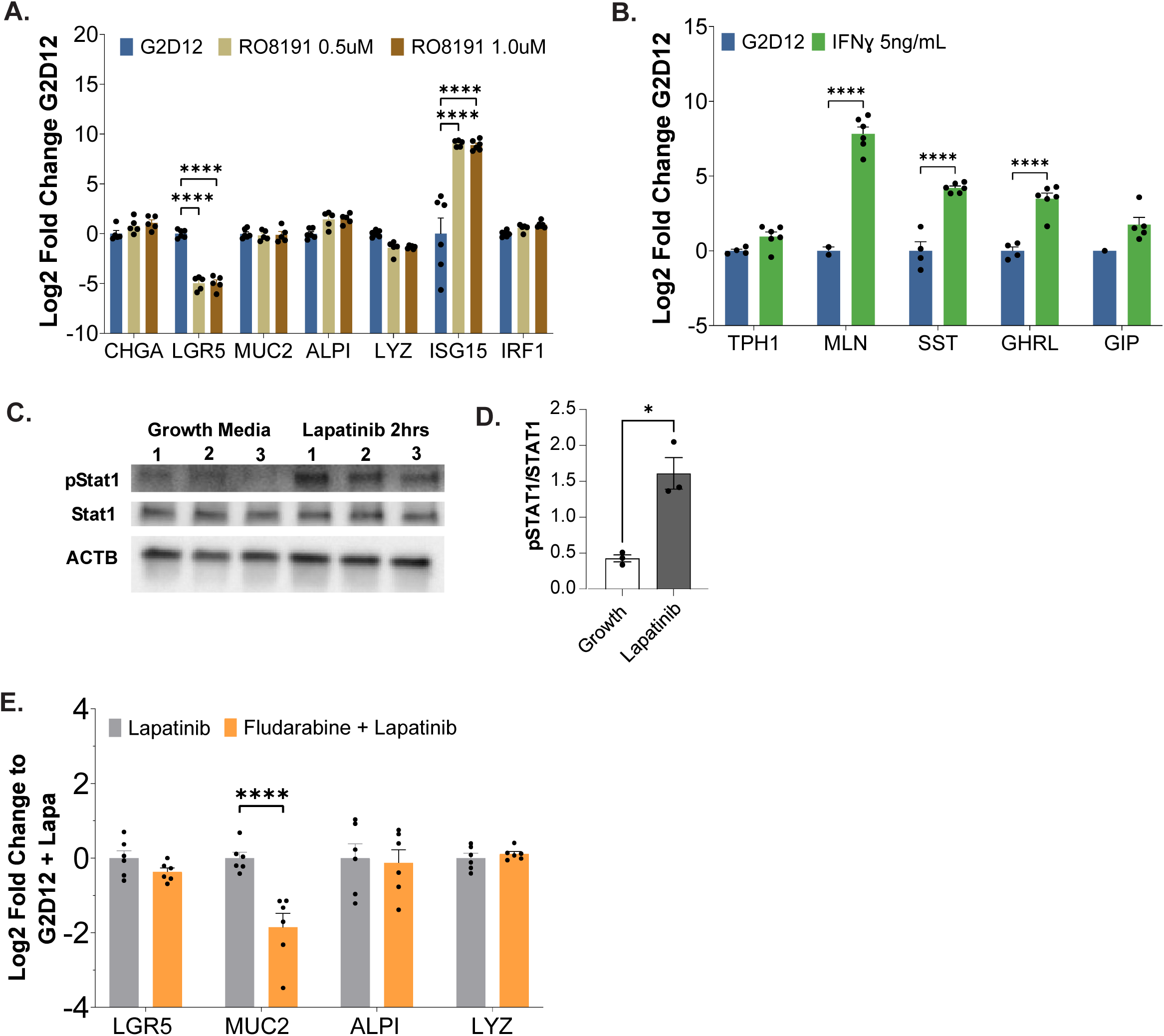
STAT1 Signaling Mediates EGFR Inhibitor–Induced EEC Differentiation in Human Organoids. (A) qPCR analysis of intestinal lineage and IFN response genes in organoids grown in G2D12 with increasing concentrations of RO8191 (type I IFN agonist), normalized to *ACTB* and plotted as log₂ fold change relative to G2D12. Representative experiment showing n = 6 wells per condition from a single organoid line. (B) qPCR analysis of EEC hormone genes in organoids grown in G2D12 with IFNγ (5 ng/mL), normalized to ACTB and plotted as log₂ fold change relative to G2D12. Representative experiment showing n = 1-6 wells per condition from a single organoid line. TPH1 = tryptophan hydroxylase 1; MLN = motilin; SST = somatostatin; GHRL = ghrelin; GIP = glucose-dependent insulinotropic peptide. (C,D) Western blot analysis (C) and quantification (D) of STAT1 phosphorylation (pSTAT1) and total STAT1 in organoids treated with lapatinib for 2 hours compared to growth media controls. Quantification is shown as the ratio of pSTAT1 to total STAT1. ACTB serves as a loading control. (E) qPCR analysis of intestinal lineage markers in organoids treated with lapatinib alone or in combination with fludarabine, normalized to *ACTB* and plotted as log₂ fold change relative to G2D12 + Lapatinib. Representative experiment showing n = 6 wells per condition from a single organoid line. LGR5 = leucine-rich repeat-containing G-protein coupled receptor 5; MUC2 = mucin 2; ALPI = intestinal alkaline phosphatase; LYZ = lysozyme. Data points are not shown when transcript levels were below the limit of detection. Bars show mean ± SEM; two-way ANOVA with Tukey correction for multiple comparisons (A, B, F); unpaired, two-tailed Student’s t-test (D). *p < 0.05; ***p < 0.001; ****p < 0.0001. Each experiment was repeated with at least three independent organoid lines.

**Supplementary Table 1. High-throughput screen compounds investigated for EEC differentiation.**

**Supplementary Table 2. Demographic information from serum samples obtained for ELISA quantification.**

**Supplementary Table 3. Growth and differentiation media components.**

**Supplementary Table 4. qPCR primers.**

**Supplementary Table 5. Reagents and Resources.**

