## Supplementary Table for "EGFR INHIBITION PROMOTES ENTEROENDOCRINE CELL DIFFERENTIATION CONTRIBUTING TO TREATMENT-ASSOCIATED DIARRHEA"

### SUPPLEMENTARY TABLES

**Supplementary Table 1. High-throughput screen compounds investigated for EEC differentiation.**

| # | Compound Name | Molar Concentration | Average %CHGA+cells/DAPI area |
|---|---|---|---|
| 1 | 10-hydroxycamptothecin | 5.5 mM | 0.0000 |
| 2 | 17- hydroxyprogesterone | 6.1 mM | 0.0000 |
| 3 | 2',3' - dideoxycytidine | 9.5 mM | 0.0013 |
| 4 | 3'-azido-3'-deoxythymidine | 7.5 mM | 0.0029 |
| 5 | 4-aminosalicylic acid | 13.1 mM | 0.0028 |
| 6 | 5-aminosalicylic acid | 13.1 mM | 0.0038 |
| 7 | 5-fluorouracil | 15.4 mM | 0.1502 |
| 8 | Abamectin | 2.3 mM | 0.2329 |
| 9 | Aceclofenac | 5.6 mM | 0.0037 |
| 10 | Acemetacin | 4.8 mM | 0.0126 |
| 11 | Acetylcholine Cl | 13.7 mM | 0.0713 |
| 12 | Acetylsalicylic acid | 11.1 mM | 0.0022 |
| 13 | Acipimox | 13 mM | 0.0252 |
| 14 | Aclarubicin | 2.5 mM | 0.5005 |
| 15 | Acycloguanosine | 8.9 mM | 0.0051 |
| 16 | Albendazole | 7.5 mM | 0.0583 |
| 17 | Alendronate | 8.1 mM | 0.0027 |
| 18 | Alfacalcidol | 5 mM | 0.0326 |
| 19 | Allopurinol | 14.7 mM | 0.0058 |
| 20 | Alprenolol HCl | 8 mM | 0.0065 |
| 21 | Alprostadil | 5.6 mM | 0.0013 |
| 22 | Altretamine | 9.5 mM | 0.0040 |
| 23 | Ambroxol | 5.3 mM | 0.0465 |
| 24 | Amfebutamone | 8.3 mM | 0.0061 |
| 25 | Amifostine | 9.3 mM | 0.2560 |
| 26 | Amiloride | 8.7 mM | 0.1754 |
| 27 | Aminophylline | 11.1 mM | 0.0095 |
| 28 | Amiodarone | 3.1 mM | 0.0120 |
| 29 | Amlodipine | 4.9 mM | 0.1648 |
| 30 | Amorolfine | 6.3 mM | 0.0092 |
| 31 | Amoxapine | 6.4 mM | 0.0117 |
| 32 | Ampicillin trihydrate | 5.7 mM | 0.0021 |
| 33 | Ampiroxicam | 4.5 mM | 0.0008 |
| 34 | Amprenavir | 4 mM | 0.0011 |
| 35 | Anagrelide | 7.8 mM | 0.0144 |
| 36 | Anastrozole | 6.8 mM | 0.0250 |
| 37 | Anethole, trans- | 13.5 mM | 0.0105 |
| 38 | Anethole-trithione (anetholtrithion) | 8.3 mM | 0.0814 |
| 39 | Aniracetam | 9.1 mM | 0.1133 |
| 40 | Apomorphine r (-) | 7.5 mM | 0.0067 |
| 41 | Apramycin | 3.7 mM | 0.0042 |
| 42 | Aprepitant | 3.7 mM | 0.0019 |
| 43 | Argatroban | 3.9 mM | 0.0027 |
| 44 | Aripiprazole | 4.5 mM | 0.0069 |

| | | | |
|---|---|---|---|
| 45 | Artemisinin | 7.1 mM | 0.0042 |
| 46 | Artesunate | 5.2 mM | 0.0058 |
| 47 | Astemizole | 4.4 mM | 0.0023 |
| 48 | Atazanavir | 2.8 mM | 0.1439 |
| 49 | Atenolol | 7.5 mM | 0.0000 |
| 50 | Atovaquone | 5.5 mM | 0.0067 |
| 51 | Atracurium besylate | 2.2 mM | 0.0023 |
| 52 | Atropine sulfate | 6.9 mM | 0.0335 |
| 53 | Auranofin | 2.9 mM | 0.1250 |
| 54 | Azaperone | 6.1 mM | 0.0020 |
| 55 | Azithromycin | 2.7 mM | 0.0011 |
| 56 | Aztreonam | 4.6 mM | 0.0020 |
| 57 | Bambuterol | 5.4 mM | 0.0161 |
| 58 | Benserazide HCl | 7.8 mM | 0.2249 |
| 59 | Benzamil | 6.3 mM | 0.0087 |
| 60 | Benzydamine | 6.5 mM | 0.0000 |
| 61 | Bepridil | 5.5 mM | 0.0039 |
| 62 | Betamethasone | 4 mM | 0.0031 |
| 63 | Betaxolol HCl | 6.5 mM | 0.1134 |
| 64 | Bexarotene | 5.7 mM | 0.0375 |
| 65 | Bezafibrate | 5.5 mM | 0.0161 |
| 66 | Bicalutamide | 4.6 mM | 0.0490 |
| 67 | Bifonazole | 6.4 mM | 0.0091 |
| 68 | Bisacodyl | 5.5 mM | 0.2536 |
| 69 | Bleomycin sulfate | 1.4 mM | 0.1924 |
| 70 | Bopindolol malonate | 5.3 mM | 0.0137 |
| 71 | Bortezomib | 5.2 mM | 0.0000 |
| 72 | Bosentan | 3.6 mM | 0.0014 |
| 73 | Bromebric acid | 7 mM | 0.0256 |
| 74 | Bromhexine HCl | 5.3 mM | 0.0118 |
| 75 | Bromocriptine mesylate | 3.1 mM | 0.2194 |
| 76 | Bumetanide | 5.5 mM | 0.1869 |
| 77 | Bupivacaine HCl | 6.9 mM | 0.0030 |
| 78 | Buspirone HCl | 5.2 mM | 0.1074 |
| 79 | Butaclamol (+) | 5.5 mM | 0.0082 |
| 80 | Butenafine | 6.3 mM | 0.0068 |
| 81 | Butyrylcholine Cl | 11.5 mM | 0.0045 |
| 82 | Cabergoline | 4.4 mM | 0.0038 |
| 83 | Calcifediol | 5 mM | 0.5787 |
| 84 | Calcipotriene | 4.8 mM | 0.0914 |
| 85 | Calcitriol | 4.8 mM | 1.6261 |
| 86 | Camptothecin | 5.7 mM | 0.0000 |
| 87 | Candesartan | 4.5 mM | 0.0058 |
| 88 | Canrenone | 5.9 mM | 0.0019 |
| 89 | Canthaxanthin | 3.5 mM | 0.0797 |
| 90 | Capsaicin | 6.5 mM | 0.6068 |

| | | | |
|---|---|---|---|
| 91 | Captopril | 9.2 mM | 0.6378 |
| 92 | Carbadox | 7.6 mM | 0.0337 |
| 93 | Carbamazepine | 8.5 mM | 0.0063 |
| 94 | Carbamyl-beta-methylcholine Cl | 12.4 mM | 0.0057 |
| 95 | Carbamylcholine Cl | 13.6 mM | 0.0130 |
| 96 | Carbidopa | 8.8 mM | 0.0340 |
| 97 | Carboplatin | 5.9 mM | 0.0062 |
| 98 | Carvedilol | 4.9 mM | 0.0322 |
| 99 | Cefepime | 4.2 mM | 0.0078 |
| 100 | Cefoperazone acid | 3.1 mM | 0.0167 |
| 101 | Cefotaxime acid | 4.4 mM | 0.0000 |
| 102 | Ceftazidime | 3.7 mM | 0.0000 |
| 103 | Celecoxib | 5.2 mM | 0.0380 |
| 104 | Cerivastatin | 4.4 mM | 0.0000 |
| 105 | Chlorambucil | 6.6 mM | 0.0033 |
| 106 | Chloramphenicol | 6.2 mM | 0.0019 |
| 107 | Chlormadinone acetate | 4.5 mM | 0.0019 |
| 108 | Chloroquine phosphate | 6.3 mM | 0.0031 |
| 109 | Chlorpheniramine maleate | 7.3 mM | 0.0011 |
| 110 | Chlorpromazine HCl | 6.3 mM | 0.0192 |
| 111 | Cilastatin | 5.6 mM | 0.0051 |
| 112 | Cilnidipine | 4.1 mM | 0.2729 |
| 113 | Cilostamide | 5.8 mM | 0.0029 |
| 114 | Cimaterol | 9.1 mM | 0.0071 |
| 115 | Cimetidine | 7.9 mM | 0.1412 |
| 116 | Cinanserin | 5.9 mM | 0.0033 |
| 117 | Ciprofloxacin | 6 mM | 0.0058 |
| 118 | Cirazoline HCl | 9.2 mM | 0.0076 |
| 119 | Citalopram | 6.2 mM | 0.0009 |
| 120 | Clarithromycin | 2.7 mM | 0.0077 |
| 121 | Clemastine fumarate | 5.8 mM | 0.0155 |
| 122 | Clenbuterol | 7.2 mM | 0.0709 |
| 123 | Climbazole | 6.8 mM | 0.0000 |
| 124 | Clinafloxacin HCl | 5.5 mM | 0.0050 |
| 125 | Clindamycin HCl | 4.7 mM | 0.0477 |
| 126 | Clindamycin palmitate | 4.7 mM | 0.0238 |
| 127 | Clindamycin PO4 | 3 mM | 0.0501 |
| 128 | Clobetasol propionate | 4.3 mM | 0.0000 |
| 129 | Clodronate disodium | 8.2 mM | 0.0367 |
| 130 | Clofarabine | 6.6 mM | 0.0000 |
| 131 | Clofibrate | 8.2 mM | 0.2565 |
| 132 | Clomiphene citrate | 4.9 mM | 0.0045 |
| 133 | Clonidine HCl | 8.7 mM | 0.0206 |
| 134 | Clopamide | 5.8 mM | 0.2319 |
| 135 | Clopidogrel sulfate | 6.2 mM | 0.0068 |
| 136 | Clopidol | 10.4 mM | 0.0040 |

| | | | |
|---|---|---|---|
| 137 | Closantel | 3 mM | 0.0043 |
| 138 | Clothiapine | 5.8 mM | 0.2612 |
| 139 | Clozapine | 6.1 mM | 0.3507 |
| 140 | Conduritol b epoxide | 12.3 mM | 0.0000 |
| 141 | Corticosterone | 5.8 mM | 0.0000 |
| 142 | Crotamiton | 9.8 mM | 0.0045 |
| 143 | Cyclocytidine HCl | 8.9 mM | 0.0168 |
| 144 | Cyclophosphamide monohydrate | 7.7 mM | 0.0106 |
| 145 | Cyclosporin a | 1.7 mM | 0.0083 |
| 146 | Cyproheptadine | 7 mM | 0.0062 |
| 147 | Cyproterone acetate | 4.8 mM | 0.0012 |
| 148 | Cytarabine | 8.2 mM | 0.0000 |
| 149 | Dacarbazine | 11 mM | 0.0148 |
| 150 | Danazol | 5.9 mM | 0.0061 |
| 151 | Daunorubicin HCl | 3.8 mM | 0.0000 |
| 152 | Debrisoquin sulfate | 11.4 mM | 0.2266 |
| 153 | Decamethonium 2Br | 7.7 mM | 0.6343 |
| 154 | Dehydroepiandrosterone | 6.9 mM | 0.0008 |
| 155 | Delavirdine mesylate | 4.4 mM | 0.0095 |
| 156 | Denbufylline | 6.2 mM | 1.0256 |
| 157 | Deprenyl HCl r (-) | 10.7 mM | 0.0691 |
| 158 | Desloratadine | 6.4 mM | 0.0018 |
| 159 | Dexamethasone | 5.1 mM | 0.0487 |
| 160 | Dextromethorphan HBr | 7.4 mM | 0.0060 |
| 161 | Diazoxide | 8.7 mM | 0.0299 |
| 162 | Dibenzepine HCl | 6.8 mM | 0.0011 |
| 163 | Diclazuril | 4.9 mM | 0.0349 |
| 164 | Diclofenac, Na | 6.8 mM | 0.0019 |
| 165 | Didanosine | 8.5 mM | 0.0125 |
| 166 | Diethylstilbestrol | 7.5 mM | 0.0040 |
| 167 | Diflunisal | 8 mM | 0.0081 |
| 168 | Dihydroergocristine mesylate | 3.3 mM | 0.0010 |
| 169 | Dihydroergotamine mesylate | 3.4 mM | 0.0136 |
| 170 | Dilazep | 3.3 mM | 0.0020 |
| 171 | Diltiazem | 4.8 mM | 0.0982 |
| 172 | Dinoprost | 5.6 mM | 0.0095 |
| 173 | Dinoprostone | 5.7 mM | 0.0069 |
| 174 | Diphenhydramine HCl | 7.8 mM | 0.0082 |
| 175 | Dipyridamole | 4 mM | 0.0831 |
| 176 | Disodium cromoglycate | 3.9 mM | 0.2191 |
| 177 | Disulfiram | 6.7 mM | 0.0010 |
| 178 | Dl-aminoglutethimide | 8.6 mM | 0.0011 |
| 179 | Dobutamine HCl | 6.6 mM | 0.0213 |
| 180 | Docebenone | 6.1 mM | 0.4194 |
| 181 | Docetaxil | 2.5 mM | 0.0859 |
| 182 | Dofetilide | 4.5 mM | 0.1474 |

| | | | |
|---|---|---|---|
| 183 | Dolasetron | 6.2 mM | 0.0138 |
| 184 | Domperidone | 4.7 mM | 0.0139 |
| 185 | Dorzolamide | 6.2 mM | 0.0202 |
| 186 | Doxazosin mesylate | 4.4 mM | 0.0033 |
| 187 | Doxifluridine | 8.1 mM | 0.0032 |
| 188 | Doxofylline | 7.5 mM | 0.0031 |
| 189 | Doxorubicin HCl | 3.7 mM | 0.0501 |
| 190 | Doxycycline HCl | 4.5 mM | 0.0018 |
| 191 | Efaroxan | 9.2 mM | 0.0173 |
| 192 | Efavirenz | 6.3 mM | 0.0012 |
| 193 | Emtricitabine | 8.1 mM | 0.1716 |
| 194 | Enalapril | 5.3 mM | 0.0000 |
| 195 | Enalaprilat | 5.7 mM | 0.0355 |
| 196 | Enoxacin | 6.2 mM | 0.0023 |
| 197 | Enrofloxacin | 5.6 mM | 0.0021 |
| 198 | Entacapone | 6.6 mM | 0.0188 |
| 199 | Epinephrine-(+)-tartrate l (-) | 11.8 mM | 0.0094 |
| 200 | Eprosartan | 4.7 mM | 0.0549 |
| 201 | Ergotamine D-tartrate | 3.4 mM | 0.0178 |
| 202 | Ergothioneine | 8.7 mM | 0.0301 |
| 203 | Erlotinib | 5.1 mM | 1.0988 |
| 204 | Escitalopram | 6.2 mM | 0.0204 |
| 205 | Esmolol | 6.8 mM | 0.0079 |
| 206 | Esomeprazole potassium | 5.8 mM | 0.0047 |
| 207 | Estradiol | 7.3 mM | 0.0021 |
| 208 | Estriol | 6.9 mM | 0.0041 |
| 209 | Estrone | 7.4 mM | 0.0033 |
| 210 | Etazolate | 6.9 mM | 0.0042 |
| 211 | Ethacrynic acid | 6.6 mM | 0.1213 |
| 212 | Ethisterone | 6.4 mM | 0.0104 |
| 213 | Etidronate 2Na | 9.8 mM | 0.0097 |
| 214 | Etoposide | 3.4 mM | 0.0219 |
| 215 | Etoricoxib | 5.6 mM | 0.0223 |
| 216 | Etretinate | 5.6 mM | 0.0055 |
| 217 | Famciclovir | 6.2 mM | 0.0010 |
| 218 | Famotidine | 5.9 mM | 0.0121 |
| 219 | Fasudil | 6.9 mM | 0.0083 |
| 220 | Felodipine | 5.2 mM | 0.4652 |
| 221 | Fenbendazole | 6.7 mM | 0.0038 |
| 222 | Fenbufen | 7.9 mM | 0.0020 |
| 223 | Fenofibrate | 5.5 mM | 0.0030 |
| 224 | Fenoldopam mesylate | 6.5 mM | 0.0032 |
| 225 | Fenoprofen | 8.3 mM | 0.0049 |
| 226 | Fenretinide | 5.1 mM | 0.3985 |
| 227 | Fentiazac | 6.1 mM | 0.0143 |
| 228 | Finasteride | 5.4 mM | 0.0027 |

| | | | |
|---|---|---|---|
| 229 | Flecainide | 4.8 mM | 0.0093 |
| 230 | Fleroxacin | 5.4 mM | 0.0000 |
| 231 | Florfenicol | 5.6 mM | 0.0027 |
| 232 | Floxuridine | 8.1 mM | 0.1154 |
| 233 | Flubendazole | 6.4 mM | 0.0416 |
| 234 | Fluconazole | 6.5 mM | 0.0000 |
| 235 | Flufenamic acid | 7.1 mM | 0.0329 |
| 236 | Flumazenil | 6.6 mM | 0.2363 |
| 237 | Flunarizine-2HCl | 4.9 mM | 0.0415 |
| 238 | Fluocinolone acetonide | 4.4 mM | 0.0129 |
| 239 | Fluoxetine HCl | 6.5 mM | 0.0048 |
| 240 | Fluperlapine | 6.5 mM | 0.0076 |
| 241 | Fluphenazine 2HCl | 4.6 mM | 0.0028 |
| 242 | Flurbiprofen | 8.2 mM | 0.0029 |
| 243 | Fluspirilene | 4.2 mM | 0.0041 |
| 244 | Flutamide | 7.2 mM | 0.0010 |
| 245 | Fluvastatin Na | 4.9 mM | 0.3082 |
| 246 | Formestane | 6.6 mM | 0.0011 |
| 247 | Fosinopril | 3.5 mM | 0.0273 |
| 248 | Ftorafur | 10 mM | 0.0042 |
| 249 | Fulvestrant | 3.3 mM | 0.0030 |
| 250 | Fumagillone | 4.4 mM | 0.2903 |
| 251 | Furafylline | 7.7 mM | 0.0650 |
| 252 | Furosemide | 6 mM | 0.0010 |
| 253 | Gabapentin | 11.7 mM | 0.0644 |
| 254 | Galanthamine HBr | 7 mM | 0.0111 |
| 255 | Gallamine triethiodide | 3.9 mM | 0.0139 |
| 256 | Ganciclovir | 7.8 mM | 0.0029 |
| 257 | Gatifloxacin | 5.3 mM | 0.0000 |
| 258 | Gefitinib | 4.5 mM | 0.7246 |
| 259 | Gemcitabine HCl | 7.6 mM | 0.0000 |
| 260 | Gemfibrozil | 8 mM | 0.0031 |
| 261 | Gentamycin sulfate | 4.2 mM | 0.0010 |
| 262 | Gestrinone | 6.5 mM | 0.0147 |
| 263 | Ginkgolide a | 4.9 mM | 0.0844 |
| 264 | Gliclazide | 6.2 mM | 0.0010 |
| 265 | Glimepiride | 4.1 mM | 0.0020 |
| 266 | Glipizide | 4.5 mM | 0.0100 |
| 267 | Glyburide | 4 mM | 0.0392 |
| 268 | Goserelin acetate | 1.6 mM | 0.0108 |
| 269 | Granisetron | 6.4 mM | 0.0327 |
| 270 | Guaiacol | 16.1 mM | 0.0086 |
| 271 | Guaifenesin | 10.1 mM | 0.0089 |
| 272 | Guanabenz acetate | 8.7 mM | 0.0389 |
| 273 | Guanfacine HCl | 7.1 mM | 0.0182 |
| 274 | Haloperidol HCl | 5.3 mM | 0.0222 |

| | | | |
|---|---|---|---|
| 275 | Harmine | 9.4 mM | 0.0119 |
| 276 | Hexestrol | 7.4 mM | 0.0038 |
| 277 | Hydrocortisone | 5.5 mM | 0.0000 |
| 278 | Hydrocortisone 21-acetate | 4.9 mM | 0.0019 |
| 279 | Hydroxytacrine maleate | 9.3 mM | 0.0084 |
| 280 | Ibandronate | 6.3 mM | 0.0570 |
| 281 | Ibudilast | 8.7 mM | 0.0981 |
| 282 | Ibuprofen | 9.7 mM | 0.1487 |
| 283 | Idarubicin | 4 mM | 0.0000 |
| 284 | Idazoxan | 9.8 mM | 0.0189 |
| 285 | Idebenone | 5.9 mM | 0.0244 |
| 286 | Idoxuridine | 5.6 mM | 0.0046 |
| 287 | Ifenprodil | 6.1 mM | 0.0724 |
| 288 | Ifosfamide | 7.7 mM | 0.0038 |
| 289 | Iloprost | 5.5 mM | 0.0161 |
| 290 | Imatinib | 4.1 mM | 0.0165 |
| 291 | Imipenem | 6.7 mM | 0.0737 |
| 292 | Imipramine HCl | 7.1 mM | 0.0056 |
| 293 | Imiquimod | 8.3 mM | 0.0010 |
| 294 | Indapamide | 5.5 mM | 0.0020 |
| 295 | Indomethacin | 5.6 mM | 0.3127 |
| 296 | Ipratropium Br | 6 mM | 0.1584 |
| 297 | Iproniazid | 11.2 mM | 0.0033 |
| 298 | Irsogladine maleate | 7.8 mM | 0.0010 |
| 299 | Isobutylmethylxanthine | 9 mM | 0.0020 |
| 300 | Isoniazid | 14.6 mM | 0.0087 |
| 301 | Isoproterenol HCl (rac) | 9.5 mM | 0.0745 |
| 302 | Itopride HCl | 5.6 mM | 0.0011 |
| 303 | Itraconazole | 2.8 mM | 0.0059 |
| 304 | Ivermectin | 2.3 mM | 0.2320 |
| 305 | Kasugamycin | 5.3 mM | 0.0504 |
| 306 | Ketanserin tartrate | 5.1 mM | 0.0023 |
| 307 | Ketoconazole | 3.8 mM | 0.1005 |
| 308 | Ketoprofen | 7.9 mM | 0.0804 |
| 309 | Ketoprofen (s) | 7.9 mM | 0.0000 |
| 310 | Ketotifen fumarate | 6.5 mM | 0.0034 |
| 311 | Lacidipine | 4.4 mM | 0.0273 |
| 312 | Lamotrigine | 7.8 mM | 0.0129 |
| 313 | Lapatinib | 3.4 mM | 3.8730 |
| 314 | Latanoprost | 4.6 mM | 0.6864 |
| 315 | Leflunomide | 7.4 mM | 0.0051 |
| 316 | Letrozole | 7 mM | 0.0231 |
| 317 | Levallorphan tartrate | 7.1 mM | 0.0202 |
| 318 | Levamisole HCl | 9.8 mM | 0.0090 |
| 319 | Levetiracetam | 11.8 mM | 0.0174 |
| 320 | Levocabastine HCl | 4.8 mM | 0.0010 |

| | | | |
|---|---|---|---|
| 321 | Levodopa | 10.1 mM | 0.0011 |
| 322 | Levofloxacin HCl | 5.5 mM | 0.0028 |
| 323 | Levonorgestrel | 6.4 mM | 0.0010 |
| 324 | Lidocaine | 8.5 mM | 0.0909 |
| 325 | Lincomycin | 4.9 mM | 0.0323 |
| 326 | Linezolid | 5.9 mM | 0.0185 |
| 327 | Lisinopril | 4.9 mM | 0.0030 |
| 328 | Lobeline | 5.9 mM | 0.1208 |
| 329 | Lofexidine | 7.7 mM | 0.0012 |
| 330 | Lomefloxacin HCl | 5.7 mM | 0.0009 |
| 331 | Lomerizine HCl | 4.3 mM | 0.0047 |
| 332 | Lomofungin | 6.4 mM | 0.0783 |
| 333 | Lomustine | 8.6 mM | 0.0449 |
| 334 | Loperamide | 4.2 mM | 0.0205 |
| 335 | Loratadine | 5.2 mM | 0.0035 |
| 336 | Lorglumide | 4.4 mM | 0.0010 |
| 337 | Losartan potassium | 4.3 mM | 0.0090 |
| 338 | Lovastatin | 4.9 mM | 0.0133 |
| 339 | L-thyroxine [(3-[4-(4-hydroxy-3,5-diiodophenoxy)-3,5-diiodophenyl]-l-alanine] | 2.6 mM | 0.0000 |
| 340 | Manidipine | 3.3 mM | 0.0202 |
| 341 | Maprotiline HCl | 7.2 mM | 0.0612 |
| 342 | Mebendazol | 6.8 mM | 0.1869 |
| 343 | Mecamylamine HCl | 12 mM | 0.1092 |
| 344 | Medroxyprogesterone 17-acetate | 16.4 mM | 0.0000 |
| 345 | Mefenamic acid | 8.3 mM | 0.0103 |
| 346 | Megestrol acetate | 5.2 mM | 0.0000 |
| 347 | Meglumine | 10.2 mM | 0.0507 |
| 348 | Melatonin | 8.6 mM | 0.0356 |
| 349 | Melengestrol acetate | 5 mM | 0.0000 |
| 350 | Meloxicam | 5.7 mM | 0.0052 |
| 351 | Melphalan | 6.6 mM | 0.0350 |
| 352 | Memantine HCl | 11.2 mM | 0.0442 |
| 353 | Mephenytoin | 9.2 mM | 0.0219 |
| 354 | Mepyramine maleate | 7 mM | 0.5200 |
| 355 | Meropenem | 5.2 mM | 0.0040 |
| 356 | Mesoridazine besylate | 5.2 mM | 0.0033 |
| 357 | Mesulergine HCl | 5.5 mM | 0.0115 |
| 358 | Metformin | 15.5 mM | 0.0106 |
| 359 | Methimazole | 17.5 mM | 0.0032 |
| 360 | Methiothepin maleate | 5.6 mM | 0.3273 |
| 361 | Methyl salicylate | 13.1 mM | 0.0373 |
| 362 | Methyldopa | 9.5 mM | 0.0013 |
| 363 | Methylprednisolone | 5.3 mM | 0.0032 |
| 364 | Methysergide | 5.7 mM | 0.0023 |
| 365 | Metoclopramide HCl | 6.7 mM | 0.0324 |

| | | | |
|---|---|---|---|
| 366 | Metoprolol tartrate | 7.5 mM | 0.0011 |
| 367 | Metronidazole | 11.7 mM | 0.0043 |
| 368 | Mevastatin | 5.1 mM | 0.0132 |
| 369 | Mianserin hcl | 7.6 mM | 0.1213 |
| 370 | Miconazole | 4.8 mM | 0.0041 |
| 371 | Mifepristone | 4.7 mM | 0.0020 |
| 372 | Miglustat | 9.1 mM | 0.0021 |
| 373 | Milrinone | 9.5 mM | 0.0000 |
| 374 | Miltefosine | 4.9 mM | 1.3260 |
| 375 | Minocycline HCl | 4.4 mM | 0.0000 |
| 376 | Minoxidil | 9.6 mM | 0.0274 |
| 377 | Misoprostol | 5.2 mM | 0.0000 |
| 378 | Mitomycin c | 6 mM | 0.0143 |
| 379 | Mitoxantrone 2HCl | 4.5 mM | 0.0000 |
| 380 | Molsidomine | 8.3 mM | 0.4383 |
| 381 | Montelukast | 3.4 mM | 0.0101 |
| 382 | Moroxydine HCl | 9.6 mM | 0.0305 |
| 383 | Myclobutanil | 6.9 mM | 0.0011 |
| 384 | Mycophenolate mofetil | 4.6 mM | 0.2204 |
| 385 | Mycophenolic acid | 6.2 mM | 0.7936 |
| 386 | Nabumetone | 8.8 mM | 0.0066 |
| 387 | Nadifloxacin | 5.5 mM | 0.0033 |
| 388 | Naftopidil 2HCl | 5.1 mM | 0.0284 |
| 389 | Nalbuphine HCl | 5.6 mM | 0.0011 |
| 390 | Naloxonazine 2HCl | 3.1 mM | 0.0024 |
| 391 | Naloxone HCl | 6.1 mM | 0.0021 |
| 392 | Naltrexone HCl | 5.9 mM | 0.0529 |
| 393 | Naltriben mesylate | 4.8 mM | 0.0025 |
| 394 | Naltrindole HCl | 4.8 mM | 0.0284 |
| 395 | Naphazoline HCl | 9.5 mM | 0.0021 |
| 396 | Naproxen | 8.7 mM | 0.3396 |
| 397 | nateglinide | 6.3 mM | 0.1305 |
| 398 | Nedaplatin | 6.6 mM | 0.0139 |
| 399 | Nefazodone | 4.3 mM | 0.0032 |
| 400 | Nelfinavir Mesylate | 3 mM | 0.0031 |
| 401 | Neomycin sulfate | 3.3 mM | 0.2402 |
| 402 | Neostigmine Br | 9 mM | 0.0146 |
| 403 | Nialamide | 6.7 mM | 0.0089 |
| 404 | Nicardipine | 4.2 mM | 0.0047 |
| 405 | Nicergoline | 4.1 mM | 0.0355 |
| 406 | Nicorandil | 9.5 mM | 0.0214 |
| 407 | Nifedipine | 5.8 mM | 0.0160 |
| 408 | Nifekalant HCl | 4.9 mM | 0.0000 |
| 409 | Niflumic acid | 7.1 mM | 0.0010 |
| 410 | Niguldipine HCl | 3.3 mM | 0.0083 |
| 411 | Nimesulide | 6.5 mM | 0.0580 |

| | | | |
|---|---|---|---|
| 412 | Nimodipine | 4.8 mM | 0.0735 |
| 413 | Nisoldipine | 5.1 mM | 0.0067 |
| 414 | Nisoxetine HCl | 7.4 mM | 0.0238 |
| 415 | Nitrendipine | 5.5 mM | 0.0180 |
| 416 | Norepinephrine-(+)-tartrate l (-) | 6.3 mM | 0.0167 |
| 417 | Norethindrone | 6.7 mM | 0.0031 |
| 418 | Norfloxacin | 6.3 mM | 0.0224 |
| 419 | Novobiocin Na | 3.3 mM | 0.1160 |
| 420 | Nystatin | 2.2 mM | 0.0035 |
| 421 | Octreotide | 2 mM | 0.0000 |
| 422 | Ofloxacin | 5.5 mM | 0.0037 |
| 423 | Olanzapine | 6.4 mM | 0.0034 |
| 424 | Olmesartan | 3.6 mM | 0.0067 |
| 425 | Olopatadine | 5.9 mM | 0.0076 |
| 426 | Oltipraz | 8.8 mM | 0.0085 |
| 427 | Omeprazole | 5.8 mM | 0.0031 |
| 428 | Ondansetron | 6.8 mM | 0.0046 |
| 429 | Oseltamivir | 6.4 mM | 0.0000 |
| 430 | Ouabain | 3.4 mM | 0.0000 |
| 431 | Oxacillin sodium monohydrate | 5 mM | 0.0010 |
| 432 | Oxaliplatin | 5 mM | 0.0989 |
| 433 | Oxatomide | 4.7 mM | 0.1301 |
| 434 | Oxcarbazepine | 7.9 mM | 0.0000 |
| 435 | Oxfendazole | 6.3 mM | 0.0058 |
| 436 | Oxibendazole | 8 mM | 0.0000 |
| 437 | Oxiconazole nitrate | 4.7 mM | 0.0011 |
| 438 | Oxotremorine sesquifumarate | 9.7 mM | 0.0150 |
| 439 | Oxymetazoline HCl | 7.7 mM | 0.0147 |
| 440 | Ozagrel | 8.8 mM | 0.0311 |
| 441 | Pamidronic acid | 8.5 mM | 0.0048 |
| 442 | Pancuronium Br | 3.5 mM | 0.1775 |
| 443 | Pantoprazole | 5.2 mM | 0.0082 |
| 444 | Pantothenic acid | 9.1 mM | 0.0009 |
| 445 | Paroxetine HCl | 6.1 mM | 0.0572 |
| 446 | Pazufloxacin | 6.3 mM | 0.0000 |
| 447 | Pefloxacine mesylate | 6 mM | 0.0024 |
| 448 | Penciclovir | 7.9 mM | 0.0022 |
| 449 | Pencillin v potassium | 5.7 mM | 0.0046 |
| 450 | Pentamidine | 5.9 mM | 0.0283 |
| 451 | Pentoxifylline | 7.2 mM | 0.0084 |
| 452 | Pergolide mesylate | 6.4 mM | 0.0161 |
| 453 | Phenamil | 6.5 mM | 0.0268 |
| 454 | Phenoxybenzamine HCl | 6.6 mM | 0.2329 |
| 455 | PhentolamineHCl | 7.1 mM | 0.0046 |
| 456 | Phenylbutazone | 6.5 mM | 0.0376 |
| 457 | Phenylpropanolamine | 13.2 mM | 0.0021 |

| | | | |
|---|---|---|---|
| 458 | Phenytoin | 7.9 mM | 0.0698 |
| 459 | Phloridzin | 4.6 mM | 0.0019 |
| 460 | Physostigmine sulfate | 7.3 mM | 0.2641 |
| 461 | Picotamide | 5.3 mM | 0.0000 |
| 462 | Pilocarpine HCl | 9.6 mM | 0.0650 |
| 463 | Pimozide | 4.3 mM | 0.0139 |
| 464 | Pinacidil | 8.2 mM | 0.0055 |
| 465 | Pindolol | 8.1 mM | 0.0166 |
| 466 | Pioglitazone | 5.6 mM | 0.0152 |
| 467 | Piperacillin | 3.9 mM | 0.0060 |
| 468 | Pirenzepine 2HCl | 5.7 mM | 0.0600 |
| 469 | Piribedil HCl | 6.7 mM | 0.0153 |
| 470 | Piroxicam | 6 mM | 0.0168 |
| 471 | Plicamycin | 1.8 mM | 0.0000 |
| 472 | Practolol | 7.5 mM | 0.0825 |
| 473 | Pramipexole | 9.5 mM | 0.0033 |
| 474 | Pranlukast | 4.2 mM | 0.0084 |
| 475 | Pranoprofen | 7.8 mM | 0.0011 |
| 476 | Pravadoline | 5.3 mM | 0.7584 |
| 477 | Pravastatin lactone | 4.9 mM | 0.0027 |
| 478 | Praziquantel | 6.4 mM | 0.0023 |
| 479 | Prazosin HCl | 5.2 mM | 0.3342 |
| 480 | Prednisolone | 5.5 mM | 0.0022 |
| 481 | Prednisone | 5.6 mM | 0.0011 |
| 482 | Pregnenolone | 6.3 mM | 0.0019 |
| 483 | Primaquine phosphate | 7.7 mM | 0.0044 |
| 484 | Procainamide | 8.5 mM | 0.0247 |
| 485 | Procarbazine HCl | 9 mM | 0.0010 |
| 486 | Procaterol hcl | 6.9 mM | 0.0316 |
| 487 | Progesterone | 6.4 mM | 0.0011 |
| 488 | Promethazine HCl | 7 mM | 0.0139 |
| 489 | Pronethalol HCl | 8.7 mM | 0.0251 |
| 490 | Propafenone | 5.9 mM | 0.0175 |
| 491 | Propofol | 11.2 mM | 0.0484 |
| 492 | Propranolol | 7.7 mM | 0.0034 |
| 493 | Propranolol HCl s(-) | 7.7 mM | 0.0689 |
| 494 | Prothionamide | 11.1 mM | 0.0043 |
| 495 | Puromycin 2HCl | 4.2 mM | 0.0000 |
| 496 | Pyrantel pamoate | 3.4 mM | 0.0012 |
| 497 | Quetiapine fumarate | 2.3 mM | 0.0097 |
| 498 | Quinacrine 2HCl dihydrate | 5 mM | 0.0010 |
| 499 | Quinapril HCl | 4.6 mM | 0.0060 |
| 500 | Quinine | 6.2 mM | 0.0090 |
| 501 | Quinpirole HCl (-)- | 9.1 mM | 0.2255 |
| 502 | Racecadotril | 5.2 mM | 0.0000 |
| 503 | Raclopride l-tartrate s(-) | 5.8 mM | 0.0147 |

| | | | |
|---|---|---|---|
| 504 | Ractopamine | 6.6 mM | 0.0069 |
| 505 | Raloxifene HCl | 4.2 mM | 0.0066 |
| 506 | Ramipril | 4.8 mM | 0.0010 |
| 507 | Ranitidine HCl | 6.4 mM | 0.0126 |
| 508 | Ranolazine 2HCl | 4.7 mM | 0.0010 |
| 509 | Rapamycin | 2.2 mM | 0.0211 |
| 510 | Rebamipide | 5.4 mM | 0.0525 |
| 511 | Remoxipride | 5.4 mM | 0.2622 |
| 512 | Retinoic acid | 6.7 mM | 0.0055 |
| 513 | Ribavirin | 8.2 mM | 0.0030 |
| 514 | Ricobendazole | 7.1 mM | 0.0092 |
| 515 | Rifampicin | 2.4 mM | 0.0626 |
| 516 | Rifamycin sv | 2.9 mM | 0.0051 |
| 517 | Rilmenidine hemifumarate | 11.1 mM | 0.0975 |
| 518 | Riluzole HCl | 8.5 mM | 0.0318 |
| 519 | Rimantadine HCl | 11.2 mM | 0.0025 |
| 520 | Risedronic acid | 7.1 mM | 0.0082 |
| 521 | Risperidone | 4.9 mM | 0.0112 |
| 522 | Rivastigmine | 8 mM | 0.0352 |
| 523 | Rocuronium bromide | 3.8 mM | 0.0000 |
| 524 | Rofecoxib | 6.4 mM | 0.0058 |
| 525 | Rolipram | 7.3 mM | 0.0021 |
| 526 | Roxatidine acetate HCl | 5.7 mM | 0.0021 |
| 527 | Roxithromycin | 2.4 mM | 0.0000 |
| 528 | Rufloxacin | 5.5 mM | 0.0023 |
| 529 | Salbutamol sulfate | 8.4 mM | 0.0501 |
| 530 | Salmeterol | 4.8 mM | 0.0078 |
| 531 | Sarafloxacin HCl | 5.2 mM | 0.0012 |
| 532 | Scopolamine HBr | 6.6 mM | 0.0000 |
| 533 | Scopolamine n-butylbromide | 5.5 mM | 0.0034 |
| 534 | Secnidazole | 10.8 mM | 0.0045 |
| 535 | Selegiline | 10.7 mM | 0.0049 |
| 536 | Sertaconazole | 4.6 mM | 0.0380 |
| 537 | Shikonin | 6.9 mM | 0.1237 |
| 538 | Sibutramine HCl | 7.1 mM | 0.0042 |
| 539 | Siguazodan | 7 mM | 0.0022 |
| 540 | Sildenafil | 4.2 mM | 0.0099 |
| 541 | Simvastatin | 4.8 mM | 0.0675 |
| 542 | Sodium phenylacetate | 12.6 mM | 0.0419 |
| 543 | Sodium phenylbutyrate | 10.7 mM | 0.0205 |
| 544 | Sotalol HCl | 7.3 mM | 0.0068 |
| 545 | Sparfloxacin | 5.1 mM | 0.0033 |
| 546 | Spectinomycin | 6 mM | 0.0240 |
| 547 | Spiperone | 5.1 mM | 0.0026 |
| 548 | Spironolactone | 4.8 mM | 0.0047 |
| 549 | Spiroxatrine | 5.3 mM | 0.0543 |

| | | | |
|---|---|---|---|
| 550 | Stanozolol | 6.1 mM | 0.0041 |
| 551 | Streptomycin sulfate | 3.4 mM | 0.0000 |
| 552 | Strychnine HCl | 6 mM | 0.0234 |
| 553 | Succinylcholine | 6.9 mM | 0.0011 |
| 554 | Sulbactam | 8.6 mM | 0.0110 |
| 555 | Sulfadiazine | 8 mM | 0.0021 |
| 556 | Sulfadimethoxine | 6.4 mM | 0.0000 |
| 557 | Sulfadoxine | 6.4 mM | 0.0055 |
| 558 | Sulfasalazine | 5 mM | 0.0000 |
| 559 | Sulindac | 5.6 mM | 0.0223 |
| 560 | Sulpiride s (-) | 5.9 mM | 0.0082 |
| 561 | Sumatriptan Succinate | 4.8 mM | 0.0050 |
| 562 | Suramin sodium | 1.5 mM | 0.0290 |
| 563 | Tacrine HCl | 10.1 mM | 0.0131 |
| 564 | Tamoxifen citrate | 5.4 mM | 0.0022 |
| 565 | Tamsulosin HCl | 4.9 mM | 0.0010 |
| 566 | Tanshinone iia | 6.8 mM | 0.0262 |
| 567 | Taurocholic acid, sodium salt hydrate | 3.9 mM | 0.0010 |
| 568 | Taxol | 2.3 mM | 0.4024 |
| 569 | Telenzepine 2HCl | 5.4 mM | 0.0443 |
| 570 | Telmisartan | 3.9 mM | 0.0031 |
| 571 | Temozolomide | 10.3 mM | 0.0000 |
| 572 | Tenatoprazole | 5.8 mM | 0.0024 |
| 573 | Tenoxicam | 5.9 mM | 0.0039 |
| 574 | Terazosin HCl | 5.2 mM | 0.0051 |
| 575 | Terbinafine HCl | 6.9 mM | 0.0167 |
| 576 | Terfenadine | 4.2 mM | 0.0043 |
| 577 | Tetracycline | 4.5 mM | 0.0041 |
| 578 | Thalidomide | 7.7 mM | 0.0000 |
| 579 | Thiamphenicol glycinate | 4.8 mM | 0.0019 |
| 580 | Thioridazine HCl | 5.4 mM | 0.0020 |
| 581 | Tibolone | 6.4 mM | 0.0010 |
| 582 | Ticlopidine HCl | 7.6 mM | 0.0192 |
| 583 | Timolol maleate (s) | 6.3 mM | 0.0684 |
| 584 | Tinidazole | 8.1 mM | 0.0000 |
| 585 | Tioconazole | 5.2 mM | 0.0000 |
| 586 | Tiotidine | 6.4 mM | 0.0327 |
| 587 | Tiotropium Br | 4.2 mM | 0.0010 |
| 588 | Tizanidine HCl | 7.9 mM | 0.1338 |
| 589 | Tobramycin (free base) | 4.3 mM | 0.0010 |
| 590 | Tolazamide | 6.4 mM | 0.0144 |
| 591 | Tolbutamide | 7.4 mM | 0.0053 |
| 592 | Tolcapone | 7.3 mM | 0.0036 |
| 593 | Tolfenamic acid | 7.6 mM | 0.0035 |
| 594 | Tolmetin Na | 7.8 mM | 0.0010 |
| 595 | Toltrazuril | 4.7 mM | 0.0029 |

| | | | |
|---|---|---|---|
| 596 | Topotecan | 4.7 mM | 0.0000 |
| 597 | Toremifene | 4.9 mM | 0.0013 |
| 598 | Tosufloxacin | 4.9 mM | 0.0045 |
| 599 | Tramadol HCl | 7.6 mM | 0.0021 |
| 600 | Tranilast | 6.1 mM | 0.1274 |
| 601 | Trans-triprolidine HCl | 7.2 mM | 0.0292 |
| 602 | Tranylcypromine | 15 mM | 0.2045 |
| 603 | Trequinsin | 4.9 mM | 0.0011 |
| 604 | Triamcinolone | 5.1 mM | 0.0035 |
| 605 | Trichloromethiazide | 5.3 mM | 0.0026 |
| 606 | Trifluoperazine | 4.9 mM | 0.2077 |
| 607 | Trifluperidol 2HCl | 4.9 mM | 0.0009 |
| 608 | Trimethoprim | 6.9 mM | 0.0032 |
| 609 | Triptorelin | 1.5 mM | 0.0053 |
| 610 | Troglitazone | 4.5 mM | 0.0096 |
| 611 | Troleandomycin | 2.5 mM | 0.0363 |
| 612 | Tropicamide | 7 mM | 0.0207 |
| 613 | Tropisetron HCl | 7 mM | 0.0013 |
| 614 | Tubocurarine Cl (+) | 3.3 mM | 0.0243 |
| 615 | Tulobuterol | 8.8 mM | 0.0064 |
| 616 | Tylosin tartrate | 2.2 mM | 0.0054 |
| 617 | Valaciclovir | 6.2 mM | 0.0029 |
| 618 | Valproic acid | 13.9 mM | 0.0735 |
| 619 | Vardenafil | 4.1 mM | 0.0187 |
| 620 | Vatalanib | 5.8 mM | 0.0121 |
| 621 | Vecuronium Br | 3.6 mM | 0.0064 |
| 622 | Venlafaxine HCl | 7.2 mM | 0.0009 |
| 623 | Verapamil | 4.4 mM | 0.1495 |
| 624 | Vidarabine | 7.5 mM | 0.0011 |
| 625 | Vinblastine sulfate | 2.5 mM | 0.0000 |
| 626 | Vincristine sulfate | 2.4 mM | 0.0000 |
| 627 | Vindesine | 2.7 mM | 0.0000 |
| 628 | Vinorelbine | 2.6 mM | 0.0000 |
| 629 | Vinpocetine | 5.7 mM | 0.0023 |
| 630 | Vitamin a (acetate) | 6.1 mM | 0.0079 |
| 631 | Xamoterol hemifumarate | 5.9 mM | 0.1216 |
| 632 | Xylazine HCl | 9.1 mM | 0.0065 |
| 633 | Yohimbine HCl | 5.6 mM | 0.0291 |
| 634 | Zafirlukast | 3.5 mM | 0.0461 |
| 635 | Zaprinast | 7.4 mM | 0.0000 |
| 636 | Zardaverine | 7.5 mM | 0.0023 |
| 637 | Zileuton | 8.5 mM | 0.0347 |
| 638 | Zoledronic acid | 7.4 mM | 0.0428 |
| 639 | zolmitriptan | 6.6 mM | 0.0192 |
| 640 | Zonisamide | 9.4 mM | 0.0746 |

**Supplementary Table 2. Demographic information from serum samples obtained for ELISA quantification.**

| Paired Controls # | Age at Dx | Gender | **Genotype** | **Treatment** |
|---|---|---|---|---|
| 1 | 37 | Female | EGFR exon 19 del | Erlotinib |
| 2 | 39 | Female | EGFR exon 19 del | Erlotinib |
| 3 | 44 | Female | EGFR exon 19 del | Erlotinib |
| 4 | 45 | Female | EGFR exon 19 del | Erlotinib |
| 5 | 45 | Female | EGFR exon 19 del | Erlotinib |
| 6 | 46 | Female | EGFR exon 19 del | Erlotinib |
| 7 | 47 | Male | EGFR L858R | Erlotinib |
| 8 | 47 | Female | EGFR exon 19 del | Erlotinib |
| 9 | 49 | Female | EGFR exon 19 del | Erlotinib |
| 10 | 49 | Female | EGFR L858R | Erlotinib |
| 11 | 50 | Female | EGFR exon 19 del | Erlotinib |
| 12 | 50 | Female | EGFR L858R | Erlotinib |
| 13 | 51 | Female | EGFR exon 19 del | Erlotinib |
| 14 | 66 | Female | EGFR L858R | Erlotinib |
| 15 | 54 | Female | EGFR exon 19 del | Erlotinib |
| 16 | 55 | Female | EGFR G719X | Erlotinib |
| 17 | 55 | Female | EGFR exon 19 del | Erlotinib |
| 18 | 56 | Female | EGFR L858R | Erlotinib |
| 19 | 56 | Female | EGFR L858R | Erlotinib |
| 20 | 58 | Male | EGFR exon 19 del | Erlotinib |
| 21 | 59 | Female | EGFR L858R | Erlotinib |
| 22 | 60 | Female | EGFR L858R | Erlotinib |
| 23 | 60 | Female | EGFR exon 19 del | Erlotinib |
| 24 | 62 | Male | EGFR L858R | Erlotinib |
| 25 | 62 | Female | EGFR exon 19 del | Erlotinib |
| 26 | 62 | Female | EGFR exon 19 del | Erlotinib |
| 27 | 63 | Female | EGFR G719X | Erlotinib |
| 28 | 64 | Male | EGFR exon 19 del | Erlotinib |
| 29 | 65 | Female | EGFR L861Q | Erlotinib |
| 30 | 65 | Female | EGFR L858R | Erlotinib |
| 31 | 67 | Female | EGFR exon 19 del | Erlotinib |
| 32 | 71 | Female | EGFR exon 19 del | Erlotinib |
| 33 | 71 | Male | EGFR exon 19 del | Erlotinib |
| 34 | 72 | Female | EGFR L858R | Erlotinib |
| 35 | 59 | Female | EGFR L858R | Erlotinib |
| 36 | 74 | Female | EGFR G719X | Erlotinib |
| 37 | 74 | Female | EGFR exon 19 del | Erlotinib |
| 38 | 77 | Female | EGFR exon 19 del | Erlotinib |
| 39 | 82 | Female | EGFR L858R | Erlotinib |
| 40 | 72 | Male | EGFR L858R | Carboplatin+Pemetrexed+Erlotinib |
| 41 | 51 | Female | EGFR exon 19 del | Erlotinib |
| 42 | 60 | Female | EGFR exon 19 del | Erlotinib |
| 43 | 57 | Female | EGFR L858R | Erlotinib |
| 44 | 62 | Female | EGFR L858R | Erlotinib |
| 45 | 62 | Female | EGFR exon 19 del | Erlotinib |
| 46 | 68 | Male | EGFR exon 19 del | Erlotinib |
| 47 | 80 | Female | EGFR exon 19 del | Erlotinib |

**Supplementary Table 3. Growth and differentiation media components**

| Growth Media | Differentiation Media |
|---|---|
| L-WRN conditioned media (65% v/v) | |
| DMEM/F12 (30% v/v) | |
| Glutamax (1% v/v) | |
| N-2 Supplement (1% v/v) | |
| B-27 Supplement (1% v/v) | |
| HEPES (10mM) | |
| Primocin (100µg/mL) | |
| Normocin (100µg/mL) | |
| A83-01 (500nM) | |
| N-Acetyl-cysteine (500µM) | |
| Recombinant Murine EGF (50ng/mL) | |
| Human [Leu15] Gastrin I (50nM) | |
| Nicotinamide (10mM) | |
| SB202190 (10µM) | |
| | DAPT (20µM) |
| | Betacellulin (20ng/mL) |
| | Tubastatin-A (10µM) |
| | PF06260933 (6µM) |
| | Tranylcypromine (1.5µM) |
| | AS1842856 (100nM) |
| | Lapatinib (1.0µM) |
| | Erlotinib (0.5µM) |
| | Tucatinib (4µM) |
| | Osimertinib (1µM) |
| | Fludarabine (5µM, 10µM) |
| | BMS-34554 (15µM) |
| | Interferon Gamma (1ng/nL, 5ng/mL) |
| | RO8191 (0.1µM, 0.5µM, 1µM) |

**Supplementary Table 4. qPCR primers**

| **Human** | | |
|---|---|---|
| **Name** | **Abbreviation** | **Identifier** |
| Beta actin | ACTB | Hs01060665_g1 |
| Intestinal alkaline phosphatase | ALPI | Hs00357579_g1 |
| Chromogranin A | CHGA | Hs00900370_m1 |
| Glucose-dependent insulinotropic polypeptide | GIP | Hs00175030_m1 |
| Ghrelin | GHRL | Hs01074053_m1 |
| Interferon alpha | IFA | Hs03044218_g1 |
| Interferon beta | IFB | Hs01077958_s1 |
| Interferon Regulatory Factor 1 | IRF1 | Hs00971965_m1 |
| Interferon Regulatory Factor 3 | IRF3 | Hs01547283_m1 |
| Interferon Regulatory Factor 9 | IRF9 | Hs00959315_m1 |
| Interferon-stimulated gene 15 | ISG15 | Hs01921425_s1 |
| Leucine-rich repeat-containing G-protein coupled receptor 5 | LGR5 | Hs00969422_m1 |
| Lysozyme | LYZ | Hs00426232_m1 |
| Motilin | MLN | Hs00757713_m1 |
| Mucin 2 | MUC2 | Hs03005103_g1 |
| Somatostatin | SST | Hs00356144_m1 |
| Signal Transducer and Activator of Transcription 1 | STAT1 | Hs01013996_m1 |
| Stimulator of Interferon Response cGAMP Interactor 1 | STING | Hs00736955_g1 |
| Tryptophan Hydroxylase 1 | TPH1 | Hs00188220_m1 |
| **Mouse** | | |
| Beta actin | ACTB | Mm02619580_g1 |
| Chromogranin A | CHGA | Mm00514341_m1 |
| Glucose-dependent insulinotropic polypeptide | GIP | Mm01254394_m1 |
| Ghrelin | GHRL | Mm00612524_m1 |
| Somatostatin | SST | Mm00436671_m1 |
| Tryptophan Hydroxylase 1 | TPH1 | Mm01202614_m1 |

**Supplementary Table 5. Reagents and Resources**

| **Reagent or Resource** | Source | Catalog Number |
|---|---|---|
| **Primary antibodies (Dilution used)** | | |
| Alexa Fluor 647-Conjugated Anti-Chromogranin A Antibody (1:100) | Novus Biologicals | NBP2-47850AF647 |
| Anti-Chromogranin A Antibody (1:100) | Agilent/Dako | M0890-2 |
| Anti-Chromogranin A Antibody (1:100) | Millipore Sigma | HPA017369-100UL |
| Anti-E Cadherin antibody (1:100) | Abcam | 1525 |
| Anti-GIP Antibody (1:100) | Invitrogen | PA5-76867 |
| Anti-Serotonin Antibody (1:100) | Abcam | ab66047 |
| Phospho-Stat1 Rabbit Antibody (1:1000) | Cell Signaling | 9167 |
| Stat1 Rabbit Antibody (1:1000) | Cell Signaling | 9172 |
| Phospho-NF-kB p65 Rabbit Antibody (1:1000) | Cell Signaling | 3033 |
| NF-kB p65 Rabbit Antibody (1:1000) | Cell Signaling | 8242 |
| Chromogranin A Polyclonal Antibody Rabbit Anti-Human/Mouse (1:300) | Proteintech | 10529-1-AP |
| Anti E-Cadherin Antibody (1:300) | Abcam | AB11512 |
| **Secondary antibodies (Dilution used)** | | |
| Donkey anti-Goat IgG (H+L) Cross-Adsorbed Secondary Antibody, Alexa Fluor 488 (1:400) | Invitrogen | A-11055 |
| Donkey anti-Mouse IgG (H+L) Cross-Adsorbed Secondary Antibody, Alexa Fluor 488 (1:400) | Invitrogen | A-21022 |
| Donkey anti-Mouse IgG (H+L) Cross-Adsorbed Secondary Antibody, Alexa Fluor 647 (1:400) | Invitrogen | A-31571 |
| Donkey anti-Rabbit IgG (H+L) Cross-Adsorbed Secondary Antibody, Alexa Fluor 488 (1:400) | Invitrogen | A-31573 |
| Donkey anti-Rabbit IgG (H+L) Cross-Adsorbed Secondary Antibody, Alexa Fluor 647 (1:400) | Invitrogen | A-21208 |
| Anti-rabbit IgG, HRP-linked Antibody | Cell Signaling | 7074 |
| **Chemicals and Enzymes** | | |
| 4′,6-diamidino-2-phenylindole (DAPI) | Thermo Fisher Scientific | D1306 |
| A83-01 | Millipore Sigma | SML0788 |
| AS1842856 | Millipore Sigma | 344355 |

| | | |
|---|---|---|
| B-27 Supplement | Thermo Fisher Scientific | 12587010 |
| Betacellulin | Peprotech | 100-50 |
| Bovine serum albumin | Millipore Sigma | 05470-1G |
| EDTA | Invitrogen | 15575020 |
| Collagenase Type I | Thermo Fisher Scientific | 17018029 |
| Diprotin A | Selleckchem | S2915 |
| Forskolin | Tocris | 6029 |
| Advanced DMEM/F12 | Thermo Fisher Scientific | 12634028 |
| DMEM GlutaMAX | Thermo Fisher Scientific | 35050079 |
| DMEM Ca2+ Free | Thermo Fisher Scientific | 21068028 |
| RNase Inhibitor | Thermo Fisher Scientific | N8080119 |
| EGF, Recombinant Murine | Peprotech | 315-09 |
| Fetal Bovine Serum Certified USA Origin | Thermo Fisher Scientific | 16000044 |
| Gastrin I (Leu15), Human | Millipore Sigma | G9145 |
| GlutaMAX | Thermo Fisher Scientific | 35050005 |
| HEPES | Thermo Fisher Scientific | 15630056 |
| Matrigel, growth factor reduced, phenol red-free | Corning | 356231 |
| N-2 Supplement | Thermo Fisher Scientific | 17502001 |
| N-Acetyl-cysteine | Millipore Sigma | A7250 |
| Nicotinamide | Millipore Sigma | N0636 |
| Paraformaldehyde, 32% | Electron Microscopy Sciences (EMS) | 15714-S |
| PF-06260933 | Millipore Sigma | PZ0272 |
| Primocin | Invivogen | ant-pm |
| Prolong Gold Antifade Mountant | Thermo Fisher Scientific | P36930 |
| Prostaglandin-E2 | Millipore Sigma | P0409 |
| SB202190 | Millipore Sigma | S7067 |
| Tranylcypromine | Millipore Sigma | P8511 |
| TRI Reagent® | Millipore Sigma | T9424-200ML |

| | | |
|---|---|---|
| Triton X-100 | Millipore Sigma | T8787-500ML |
| TruStain FcX™ (Fc Receptor Blocking Solution), Human | Biolegend | 422301 |
| TrypLE Express | Thermo Fisher Scientific | 12605010 |
| Tubastatin-A | Selleckchem | S8049 |
| Y-27632 dihydrochloride | Tocris | 1254 |
| Lapatinib | Selleckchem | S1028 |
| Erlotinib | Selleckchem | S1023 |
| Tucatinib | Selleckchem | S8362 |
| Osimertinib | Selleckchem | S7297 |
| Fludarabine | Selleckchem | S1491 |
| BMS-34554 | Selleckchem | S8044 |
| Interferon Gamma (IFN- $\gamma$) | MedChem Express | HY-P7025 |
| RO8191 | MedChem Express | HY-W063968 |
| RIPA Lysis and Extraction Buffer | Thermo Fisher Scientific | 89900 |
| Pierce™ BCA Protein Assay Kits | Thermo Fisher Scientific | 23227 |
| Trans-Blot Turbo 5X Tranfer Buffer | BioRad | 10026938 |
| Trans-Blot Turbo RTA Mini 0.2 µm PVDF Transfer Kit, for 40 blots | BioRad | 1704272 |
| Bovine Serum Albumin | Sigma | A9418-50G |
| SignalFire™ ECL Reagent | Cell Signaling | 6883P3 |
| **Commercial Assays** | | |
| Direct-zol RNA Microprep Kit | Zymo Research | R2061 |
| High-Capacity cDNA Reverse Transcription Kit | Thermo Fisher Scientific | 4368813 |
| TaqMan™ Universal PCR Master Mix | Thermo Fisher Scientific | 4304437 |
| Human GIP (total) ELISA Kit | Millipore Sigma | EZHGP1-S4K |
| Human Motilin (total) ELISA Kit | Novus | NBP2-66719 |
| Human 5-HT (total) ELISA Kit | Abcam | ab133053 |
| Human Somatostatin (total) ELISA Kit | Novus | NBP2-80269 |
| Mouse 5-HT ELISA Kit | Immusmol | BA-E-5900R |
| U-PLEX Custom Metabolic Group 1 (ms) Assays; GLP-1, PYY | Meso Scale Discovery | K152ACM |
